# Charting the Expansive Outer Membrane β-Barrel Landscape in *Pseudomonas aeruginosa*

**DOI:** 10.64898/2026.11.17.767137

**Authors:** Elina Karimullina, Lukasz Jaroszewski, Joshua Tourigny, Dhruvin Patel, Yirui Guo, Joe J. Harrison, Tatiana Rogasevskaia, Dominika Borek, Adam Godzik, Alexei Savchenko

## Abstract

Outer-membrane β-barrel (OMBB) proteins are central to Gram-negative bacterial physiology, mediating nutrient acquisition and secretion while shaping host interactions and antimicrobial permeability. Yet tools to catalog the diversity and functional potential of the OMBB proteome in these bacteria remain incompletely defined. Here, we combine proteome-scale structure prediction analysis and experimental validation to systematically map the OMBB proteome of the opportunistic pathogen *Pseudomonas aeruginosa*. We identify 105 proteins containing predicted OMBB domains with 8–26 β-strands, substantially expanding the repertoire across major structural classes and revealing previously unrecognised domain architectures. Comparison with experimentally defined structures shows that prediction accurately captures global barrel architecture but systematically overestimates β-strand continuity and underrepresents flexible extracellular loops, key determinants of molecular recognition. We then defined the cryo-EM structures of PA2590 and PA3535 (EprS), confirming their predicted functions as a TonB-dependent transporter and an autotransporter, respectively. Using structural prediction, we also identified the phospholipase A1-like activity in the uncharacterized autotransporter PA4579, linking a predicted catalytic site to function. Together, our findings expand the structural and functional landscape of the *P. aeruginosa* OMBB proteome and establish a generalizable framework for converting proteome-scale structural predictions into experimentally testable mechanisms, enabling discovery of functions hidden within poorly annotated outer-membrane proteomes.

## INTRODUCTION

*Pseudomonas aeruginosa* is a leading cause of healthcare-associated infections worldwide and remains a high-priority antimicrobial-resistant pathogen for global surveillance and therapeutic development^1,2^. This Gram-negative bacterium is known to adapt to a wide variety of environmental habitats^3,4^, withstand immune responses through adhesion, and biofilm formation, and is capable of residing in virtually all host tissues^4,5^. Individuals with a weakened immune system, as well as those with disruption of normal physiologic functions caused by cancer, severe burns, or chronic pulmonary disorders, are particularly vulnerable to *P. aeruginosa* infections^1^, while multidrug resistance complicates treatments^6,7^.

Research on *P. aeruginosa*’s pathogenesis shows a multifactorial process driven by characteristics of *P. aeruginosa*’s cell envelope^4,5^. In particular, a number of outer membrane proteins (OMPs) are necessary for successful infections in eukaryotic hosts and maintaining antibiotic resistance^8^. As in other Gram-negative bacteria, *P. aeruginosa*’s outer membrane (OM) is an asymmetric lipid bilayer composed of lipopolysaccharides (LPS) in the outer leaflet and phospholipids in the inner leaflet. Interactions between LPS molecules and divalent cations stabilize the membrane and create a permeability barrier that restricts the passive diffusion of many compounds. Transport across the OM therefore depends on specialized OMPs.

A major structural class of OMPs contains an outer membrane β-barrel (OMBB) domain. In these proteins, antiparallel β-strands form a membrane-spanning cylindrical barrel, with the number and arrangement of strands defining its overall dimensions. The barrel’s exterior is hydrophobic and interacts with the lipid bilayer, whereas lumen-facing residues and internal structural elements, including constriction loops and plug domains, define the effective pore size and chemical environment. In channel-forming OMBBs, these features determine selectivity for solutes and ions based on their size, charge, and other physicochemical properties^9,10^. Water molecules within the lumen provide an aqueous pathway across the hydrophobic membrane and can also influence the pore’s electrostatic and solvation properties^9^. Because many OMBB proteins function as diffusion channels, the terms “OMBB protein” and “porin” have sometimes been used interchangeably^11^. However, not all OMBB proteins are porins, and it is now recognized that members of this structural class also contribute to cell-envelope stability, signal transduction, adhesion, and immune evasion^9,12^.

Due to their essential role in pathogenesis and antibiotic resistance, *P. aeruginosa* OMBB proteins have been a focus of intense research^13–15^, including genomic analysis^16–21^, proteomics studies^22,23^, and individual molecular function characterization^24–29^. Collectively, these studies indicate that *P. aeruginosa* relies predominantly on substrate-selective OM channels and lacks an abundant, high-conductance general diffusion porin comparable to OmpF or OmpC in *Enterobacterales*^9,25^. OprF, encoded by *oprF* (*PA1777*), is the most abundant non-lipoprotein OM protein in *P. aeruginosa* and an OmpA-family homolog, rather than a structurally established general porin^13^. *P. aeruginosa* also encodes close to twenty OMBB proteins from the OprD (Occ) family defined by 18-stranded β-barrels, and over thirty TonB-dependent transporters (TBDT), which contain 22-stranded β-barrels^13,24^. The proton motive force (PMF) transmitted from the TonB−ExbB−ExbD complex (TonB complex) located in the inner membrane is used by TBDTs to transport specific compounds, including ion-chelating siderophores, heme, organic sulfur, and certain aromatics^30–33^. In addition to these two large groups, individual smaller *P. aeruginosa* OMBB proteins with only 8 β-strand barrel - OprH and OprG have been characterized^34–36^. A previous global study into the role of specialized *P. aeruginosa* OMBB proteins in the uptake of nutrients and antibiotics estimated several dozen different OMBB proteins^8^; however, the full account of *P. aeruginosa* OM proteome is lacking, with a significant number of OMPs remaining cryptic.

Recent advances in the accuracy and scalability of 3D protein structure prediction^37,38^ provide an opportunity to more accurately estimate the repertoire of *P. aeruginosa* OMBB proteins. By analysing structural models for the complete *P. aeruginosa* proteome, we identified more than 100 proteins predicted to contain OMBB domains comprising 8 to 26 β-strands (Supplementary Table 1). This repertoire encompasses all previously characterized or predicted *P. aeruginosa* OMBB proteins while substantially expanding the diversity of known OMBB classes, including the extensively studied OprD (Occ) family. Beyond increasing the apparent scale of the OMBB proteome, our analysis identified previously unrecognized domain architectures and accessory structural elements that will help to close substantial gaps in our understanding of OMBB-mediated transport and host interactions.

To experimentally validate this expanded repertoire, we determined the cryo-EM structures of two cryptic OMPs: PA3535 and PA2590, representing 12- and 22-stranded OMBB architectures, respectively. We further demonstrated the enzymatic activity and identified catalytically important residues in the previously uncharacterized autotransporter PA4579, consistent with structural predictions of its passenger domain.

Together, these results establish the structural catalogue of the *P. aeruginosa* OMBB proteome, defining its scale, architectural diversity, and functional potential. The resulting catalogue of β-strand topology, domain organization, and surface-exposed loops provides a framework for generating testable hypotheses concerning substrate recognition, pore gating, antibiotic permeability, and host interactions, while identifying accessible epitopes for immunological targeting. More broadly, our study demonstrates how proteome-scale, AI-driven high-quality structure prediction coupled with targeted experimental validation can transform poorly annotated proteomes into mechanistically informative maps. In this case, it provided a roadmap for dissecting transport, virulence, and multidrug-resistance mechanisms in *P. aeruginosa*.

## Results

### Proteome-wide identification of OMBB proteins in *P. aeruginosa* using multipronged *in silico* analysis

To assess the number of outer membrane β-barrel (OMBB) domains, we used the *Pseudomonas aeruginosa* PAO1 genome (taxid 208964) and predicted protein sequences from the UniProt reference proteome database (UniProt proteome: UP000002438, genome assembly GCA_000006765.1)^39^. Next, we modelled *P. aeruginosa* PAO1 proteome (5563 proteins in total) using the latest version of AlphaFold (AF) algorithm, AlphaFold3 available on the AlphaFold server^40^, and installed locally, and analysed it for the presence of predicted OMBB domains. In addition, for some proteins, alternative models have been calculated by other structure prediction algorithms, including ESM-Fold^41^.

We identified proteins predicted to contain the OMBB domain in the *P. aeruginosa* proteome using a cascade of methods, starting with a close-sequence-similarity search with BLASTP^42^, followed by remote homology detection with HHPred^43^, and analysis of structural models built with AF using FATCAT structural alignment program^44^. A search using BLASTP identified 81 proteins with close sequence similarity to ECOD database entries classified as OMBBs (outer membrane meander beta-barrel domain ECOD topology). At the same time, the sensitive remote homology detection method HHPred identified 100 proteins with significant sequence similarity to this ECOD topology. This prediction included all proteins identified by BLASTP while suggesting 19 additional proteins containing a domain with this topology. Structural analysis of predicted structural models using FATCAT identified 105 proteins with significant similarity to OMBBs from the ECOD database. This subset included all proteins identified with BLASTP and HHPred. After individual verification, the final list included 105 proteins in *P. aeruginosa* proteome that were confirmed to contain a predicted OMBB domain. All identified proteins were predicted to contain only a single OMBB domain, so the total number of identified OMBB domains (105) was equal to the total number of predicted OMBB domain-containing proteins.

### Classification and Functional Domain Organization of predicted *P. aeruginosa* OMBB Proteome

Analysis of the 105 *P. aeruginosa* proteins predicted to contain outer-membrane β-barrel (OMBB) domains revealed considerable structural and functional diversity (Supplemental Table SX). The largest class comprised 35 proteins with predicted 22-stranded OMBB and N-terminal plug domains, the characteristic architecture of TonB-dependent transporters (TBDTs; TCDB family 1.B.14)^31^. The next largest group consisted of 23 predicted 18-stranded OMBB reminiscent predominantly of Opr-family substrate-selective channels (TCDB family 1.B.25), together with the AlgE alginate exporter (TCDB family 1.B.13) and three additional predicted OMBBs lacking confident family-level functional assignments. The 17 proteins contained predicted 16-stranded OMBB similar to anion-selective OprP/OprO porins and carbohydrate-selective OprB/OprB2 porins (TCDB families 1.B.5 and 1.B.19, respectively), as well as Omp85/TpsB-superfamily proteins involved in outer-membrane protein assembly or secretion. The latter included BamA and TamA (TCDB family 1.B.33), TpsB secretion channels (TCDB family 1.B.20), and the Omp85-related protein PlpD. This class also contained the PelB exopolysaccharide exporter and several uncharacterized proteins. The 17 predicted 12-stranded OMBBs included three characterized or predicted type Va autotransporters (TCDB family 1.B.12), five FapF-family channels associated with functional amyloid secretion or putative hydrophobic-compound uptake, two Tsx-like nucleoside-channel candidates (TCDB family 1.B.10), a PcoB-family protein implicated in copper homeostasis, a putative Mg²⁺-uptake-associated channel, and two functionally uncharacterized channel candidates.

The remaining proteins with predicted OMBB domains represented smaller structural classes, with five of them predicted to contain 8-stranded β-barrels: the OmpA-like structural protein OprF (TCDB family 1.B.6), OprG of the OmpW family (TCDB family 1.B.39), the outer-membrane-stabilizing protein OprH, and the lipid A-modifying enzymes PagP and PagL. Four proteins were predicted to contain 24-stranded OMBBs reminiscent of fimbrial usher proteins (TCDB family 1.B.11). Three proteins contained 14-stranded OMBBs similar to those characterized in FadL-family channels for fatty acids and other hydrophobic substrates (TCDB family 1.B.9). We also identified one protein, PA0595, with predicted single 26-stranded OMBB similar to LptD-family LPS exporter (TCDB family 1.B.42).

Based on our annotation analysis of predicted OMBB-containing proteins, 70 of the 105 (67%) lacked direct experimental characterization at the individual-protein level. Reassessment of the literature during manuscript preparation identified prior direct functional evidence for PA4661/PagL^45,46^ and PA0328/AaaA^47^, reducing the number of experimentally uncharacterised proteins with predicted OMBB domains to 68 (65%).

To evaluate the reliability of OMBB domain predictions, we compared the identified proteins with experimentally characterized structures available in the Protein Data Bank (PDB). At the time of our analysis, 35 of the 105 predicted OMBB proteins had been at least partially structurally characterized. Six additional proteins shared high sequence identity (>=50%) with structurally characterized homologues, while another 39 exhibited statistically significant sequence similarity (BLASTP E-value ≤0.05) to proteins with experimentally determined structures, further supporting the presence of OMBB domains. The complete classification is available at https://proteins.pathogens3d.org/dataset/pseudomonas-porins. Because AF3 generates three-dimensional structural predictions without requiring a closely related experimentally determined structural homologue, we next assessed how well AF3 models recapitulate experimentally observed OMBB architectures. To this end, we systematically compared AF3-generated models with experimentally determined structures of *P. aeruginosa* OMBB proteins. This analysis revealed that AF3 models generally exhibited more regular β-barrel architectures than their experimentally defined counterparts, including more uniform β-strand geometries and hydrogen-bonding networks. This trend was further supported by comparisons with the *de novo* structures experimentally determined in this study, as discussed below.

We next examined the domain architecture of all predicted OMBB-containing proteins using the ECOD-based hierarchical annotation workflow. Domain assignments were first obtained by sequence similarity searches using BLASTP^48^, followed by remote homology detection with HHpred^43^ and structural similarity analysis of AF3 models using FATCAT^49,50^. Progressive application of these complementary approaches substantially increased annotation coverage, assigning 159 ECOD topology-classified domains by BLASTP, 174 by HHpred, and 201 by FATCAT (Fig. 1).

**Figure 1.**
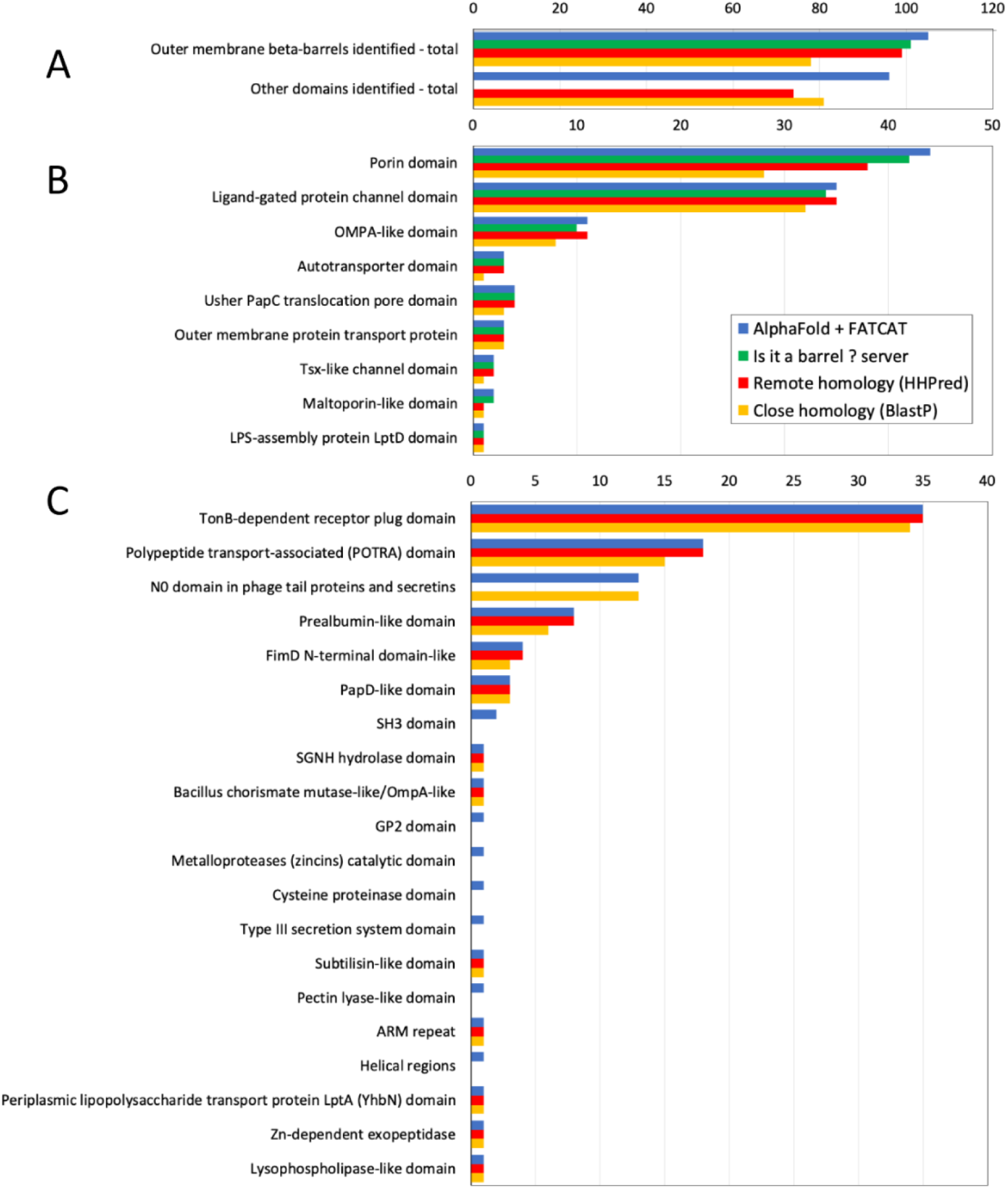
The number of domains identified in OMBB proteins from *P. aeruginosa* by different *in silico* methods. The number of domains recognized by close homology (BLASTP), remote homology (HHPred), and structural similarity of AlphaFold3 models as detected by FATCAT and IsItABarrel program, are shown as yellow, red, blue, and green bars, respectively (IsItABarrel program is only applicable to beta-barrel domains). A) the total number of outer membrane meander beta-barrel domains and the total of other domains identified in *P. aeruginosa* proteome. B) the number of outer membrane meander beta-barrel domains identified by each method divided into domain types (based on ECOD database topologies). C) the number of other domains accompanying OMMBBs identified by each method divided into domain types (based on ECOD database topologies).

Among OMBB domains, porin ECOD topology represented the largest group, occurring in 43 proteins (with 21, 17, and 5 domains consisting of 18, 16, and 12 strands, respectively), followed by Ligand-gated protein channel domain ECOD topology (corresponding to TBDTs, consisting of 22 strands each), identified in 35 proteins. Domains with OmpA-like ECOD topology were predicted in 12 proteins (7 and 5 barrels containing 12 and 8 strands, respectively), whereas all remaining OMBB classes collectively accounted for 15 proteins.

The most common accessory domain was the TonB-dependent receptor plug ECOD domain, with one copy present in all 35 predicted 22-strand OMBBs. Other frequently occurring accessory ECOD topologies included the polypeptide transport-associated (POTRA) domain (18 occurrences in 8 proteins), the N0 domain found in secretins and phage tail proteins (13 occurrences in 13 proteins), and the prealbumin-like domain (8 occurrences in 4 proteins). Sixteen additional accessory domain types together accounted for the remaining 22 annotated domains.

The correspondence between β-strand number, ECOD topology, and TCDB classification was unambiguous for 22-stranded OMBBs corresponding to TonB-dependent transporters, all of which were assigned to the ECOD ligand-gated protein channel topology and TCDB family 1.B.14. Similarly, all 24-stranded β-barrels were classified as Usher/PapC translocation pores in ECOD and belonged to TCDB family 1.B.11, whereas all 14-stranded predicted OMBBs were assigned to the Outer membrane protein transport topology in ECOD and TCDB family 1.B.9. In contrast, OMBBs belonging to other ECOD topologies showed less consistent correspondence with individual TCDB families, precluding a straightforward one-to-one mapping between structural topology and TCDB classification.

Domain combinations in predicted OMBB containing proteins exhibited clear functional organization rather than random associations. Porin ECOD domain topology occurred either alone or in combination with one or more POTRA domains, whereas POTRA domains were observed exclusively in porin-containing proteins. Likewise, ligand-gated protein channel domains (the hallmark of TonB-dependent transporters) were almost invariably associated with either the TonB-dependent receptor plug ECOD domain or the N0 domain of secretins and phage tail proteins, neither of which was detected in association with other OMBB classes. In contrast, OmpA-like domains were typically found as isolated structural modules (https://proteins.pathogens3d.org/protein/PA2590/pseudomonas-porins).

Our analysis suggested that the sequence and structure diversity is the largest in 22-and 12-strands groups of predicted OMBBs, while the database annotations of these groups in UniProt and <u>Pseudomonas.com</u> databases^18^ are least developed. Specifically, 57% of 22-strand OMBBs revealed by our analysis are annotated in UniProt as hypothetical proteins or with a generic “Probable TonB-dependent receptor,” and the same percentage for 12-strand OMBBs is 40% -as compared to less than 30% for all the remaining OMBBs. Similar percentages in both groups have no statistically significant BLAST hits to any structurally characterized protein in the PDB. While some of the proteins in these groups have been identified in literature as transmembrane barrels, our analysis is, at this point, the most comprehensive modeling and analysis effort for these groups of OMBBs. Thus, we focused on analysis of representatives of these two OMBB groups.

### *P. aeruginosa* TonB-dependent transporters (TBDTs) have substantial structural and functional diversity

Our analysis identified 22-stranded OMBB proteins as the largest group within the *P. aeruginosa* OM proteome. These OMBBs are the hallmark of TonB-dependent transporters (TBDTs), which mediate proton-motive-force (PMF)-dependent acquisition of essential ions and nutrients^31,33^. At the time of our analysis, the *Pseudomonas* genome database (https://pseudomonas.com/) provided annotations for only 11 TBDTs in *P. aeruginosa*. Because many predicted TBDTs remain incompletely curated, we manually reviewed the literature and annotated all 22-stranded OMBB proteins identified in our analysis. Evidence for a functional, regulatory, or phenotypic role was available for 27 of 35 proteins (Supplementary Table 1), although the physiological ligands of many of these receptors remain unknown. Experimentally supported nutrient-uptake functions include ferrisiderophore receptors PA2398/FpvA, PA4221/FptA, PA2466/FoxA, PA0470/FiuA, PA1910/FemA, PA2688/PfeA, PA0931/PirA, PA4168/FpvB, PA4675/ChtA, PA3268/ActA, and PA4156/FvbA; the ferric-citrate receptor PA3901/FecA; the heme receptors PA4710/PhuR and PA3408/HasR; the pseudopaline receptor PA4837/CntO, which mediates Zn and Ni acquisition; and the copper transporter PA3790/OprC. PA1302/HxuA functions primarily in heme sensing and cell-surface signalling, with a lesser contribution to heme acquisition. Other assignments remain provisional. PA1271 is predicted by homology to be a BtuB-like cobalamin receptor, although vitamin B12 transport has not been demonstrated experimentally in *P. aeruginosa*. Transcription of PA0781/ZnuD-like, PA1922, and PA2911 is induced under zinc limitation, consistent with possible roles in zinc homeostasis, but receptor-specific zinc transport has not been established. PA0434/CrtA is copper responsive and is exploited by S-type pyocins SX1 and SX2 for outer-membrane translocation^51^. Subsequent family-level analysis placed PA0434 within a group of CrtA-related TonB-dependent copper transporters, although its physiological substrate remains unconfirmed. Likewise, the physiological ligands of PA2590, PA2089, and PA2289 remain unknown.

Cross-referencing the predicted 22-stranded OMBB proteins against experimentally determined structures in the Protein Data Bank (PDB; https://www.rcsb.org/) revealed that only nine had been structurally characterized at the time of analysis (Supplementary Table 2). Eight additional proteins were confidently predicted to adopt the characteristic TBDT architecture but remain annotated as proteins of unknown function, without experimentally defined ligand specificity or structural information. While this manuscript was in preparation, structures of three further members (PA1910, PA3063 and PA3648) became available (Supplementary Table 2). Thus, despite the established importance and apparent functional diversity of the *P. aeruginosa* TBDT repertoire, its structural and mechanistic characterization remains incomplete. This knowledge gap is particularly important because extracellular loop architecture, which varies substantially across TBDTs, is a major determinant of substrate recognition and uptake^33^. Structural characterization of the understudied TBDTs therefore provides an important route to identifying new substrate specificities and understanding nutrient acquisition under the nutrient-limited conditions encountered during infection.

Part of poorly characterised TBDTs, PA2590 was originally annotated as a conserved hypothetical protein in the *P. aeruginosa* PAO1 proteome and remains so in major reference databases, including NCBI RefSeq (NP_251280.1) and KEGG. Comparative modelling resources (see https://proteins.pathogens3d.org/protein/PA2590/pseudomonas-porins) subsequently classified PA2590 as a TonB-dependent β-barrel transporter based on primary sequence similarity. A recent host–pathogen spatial transcriptomics study in a murine model of *P. aeruginosa* ocular infection identified PA2590 as a major virulence determinant, with disruption of the corresponding gene significantly attenuating infection^52^. Although this study suggested a possible role for PA2590 in iron and/or cobalamin (vitamin B12) uptake, these assignments remained speculative in the absence of structural or mechanistic characterization.

To define the molecular architecture of PA2590, we determined its structure by single-particle cryo-electron microscopy (cryo-EM) to an overall resolution of 2.93 Å (Supplementary Fig. 1, A– D). Consistent with structural prediction, PA2590 adopts the canonical two-domain architecture of a TBDT, comprising a C-terminal 22-stranded antiparallel β-barrel (residues 278–883) and an N-terminal globular plug domain (residues 134–270) that occludes the barrel lumen (Fig. 2). No interpretable density was observed for the N-terminal region preceding the plug domain, which is predicted by the Pathogens3D database (https://proteins.pathogens3d.org/protein/PA2590/pseudomonas-porins) to form a TonB-related domain. The extracellular surface of the β-barrel is capped by extended loops that partially shield the plug domain and form a shallow, surface-exposed binding pocket. In contrast, the periplasmic face consists primarily of short turns connecting adjacent β-strands, also characteristic of TBDTs (Fig. 2).

**Figure 2.**
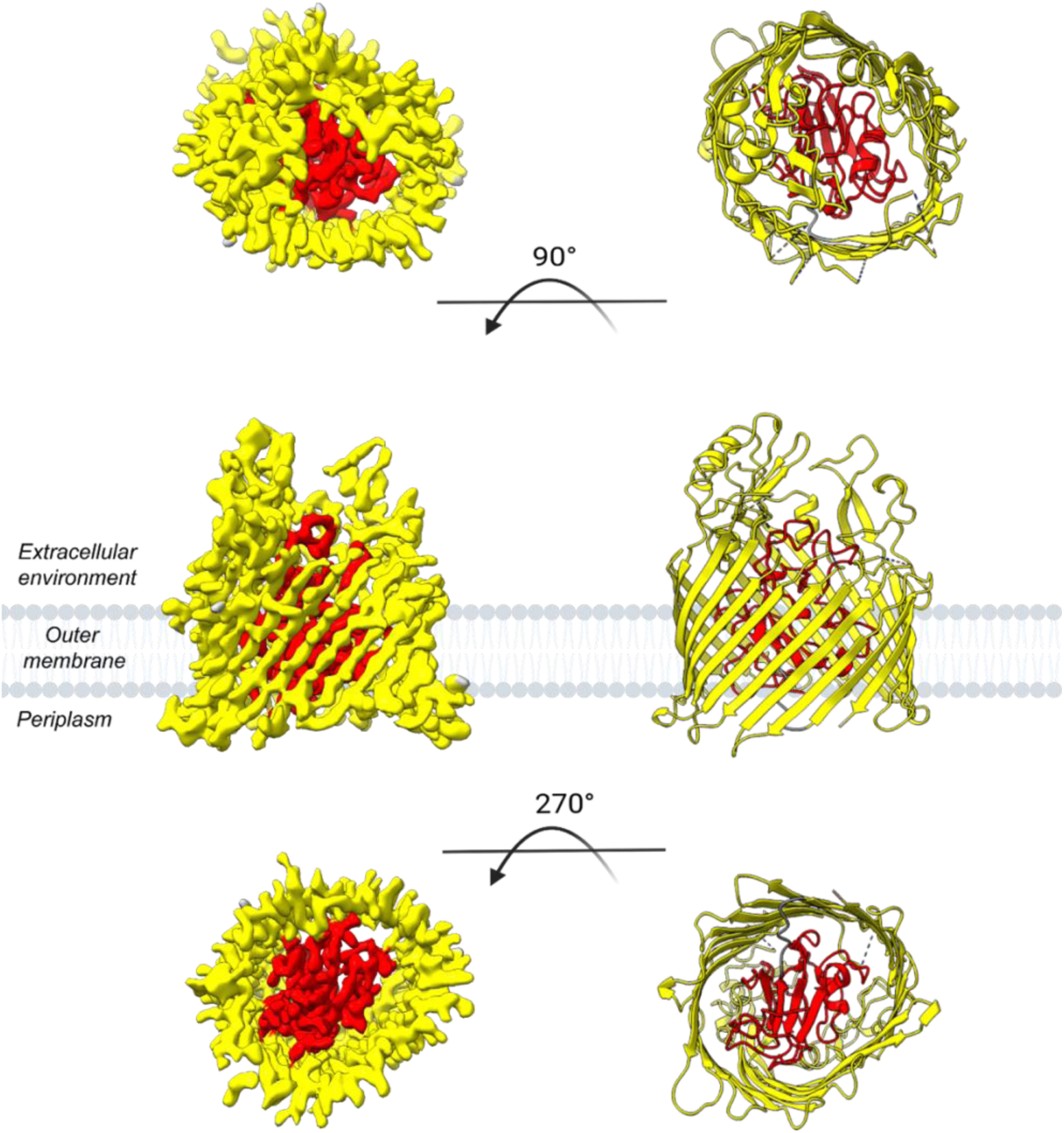
Cryo-EM structure of TonB-dependent transporter of unknown function (PA2590). Cartoon representations (right) and corresponding density maps (left) are shown in three orientations: extracellular view (top), side view within the outer-membrane plane (middle), and periplasmic view (bottom). PA2590 comprises a 22-stranded antiparallel β-barrel forming a closed outer-membrane barrel, with an N-terminal plug/cork domain positioned inside the barrel lumen. The plug domain is shown in red ( residues 134–270), the β-barrel domain is shown in yellow (residues 278–883), and the linker region between the plug and barrel corresponds to residues 271–277. The outer-membrane plane is indicated schematically in gray.

The extracellular loops of the PA2590 OMBB contain both α-helical and β-strand insertions that frame the entrance to the ligand-transport pathway (Fig. 3A). Density for the C-terminal extracellular loops L6–L11 was sufficiently well resolved to define their approximate boundaries and structural features. L6 is a short connector spanning residues 560–563, whereas L7 (592–631), L8 (662–697), L9 (742–765), L10 (789–816), and L11 (840–874) form larger surface-exposed loops that shape the extracellular opening of the barrel. Several of these loops contain short secondary-structure elements, including α-helices and α-turns in L7, L8, L9, and L11 and β-strand or hairpin-like elements in L7 and L10, respectively. In contrast, density corresponding to L2 (325–344), L3 (377–402), L4 (436–459), and L5 (505–526) was not resolved, most likely because of substantial flexibility in these regions (Fig. 3B).

**Figure 3.**
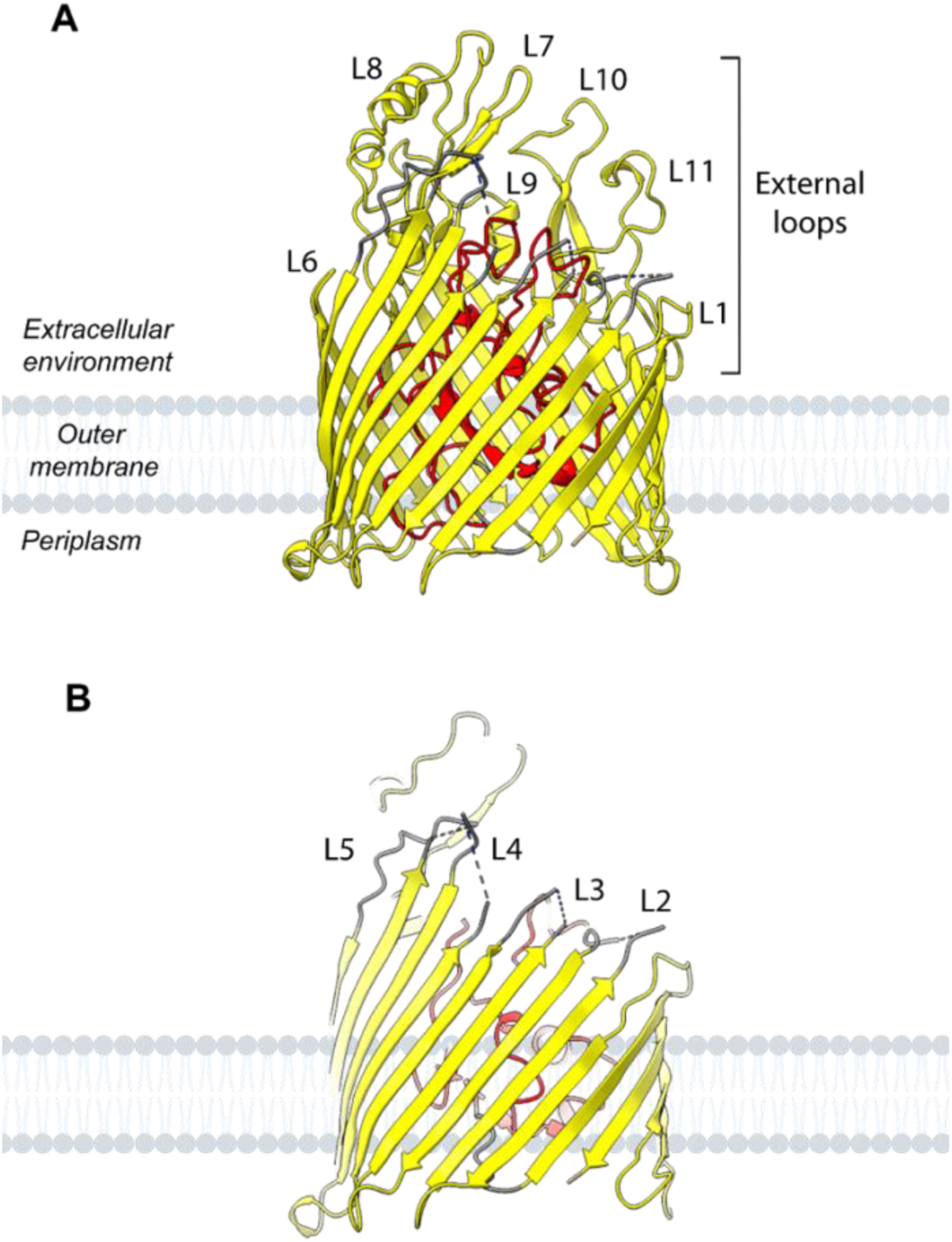
Extracellular loop architecture of PA2590. (A) Side-view cartoon representation of PA2590 highlighting the core β-barrel in yellow and the plug scaffold in red, with the positions of the extracellular loops indicated. Extracellular loops containing poorly resolved or unresolved regions in the cryo-EM density are shown in grey. (B) The same orientation with the posterior portion of the barrel removed, revealing the regions corresponding to poorly resolved extracellular loops in the cryo-EM density and illustrating their pronounced flexibility.

Superposition of the experimental PA2590 structure with its AF3-derived model showed close agreement in the core β-barrel and plug scaffolds (RMSD, 0.716 Å over 575 pruned atom pairs), confirming the overall accuracy of the predicted architecture. However, substantially greater deviation was observed across the complete structures (RMSD, 2.071 Å over 672 atom pairs), particularly in extracellular loops and periplasmic turns (Fig. 4). Several β-strands were also predicted to extend beyond their experimentally observed boundaries. For example, β-strand 10 comprises residues Ser530–Thr542 in the experimental structure compared with Gly527–Thr545 in the predicted model; β-strand 4 spans Phe346–Phe357 experimentally versus Ala345–Leu359 in the model; and β-strand 2 comprises Gly297–Ala305 experimentally versus Thr295–Phe306 in the model. Similar differences were observed in extracellular loops L7 and L8. L7, connecting β-strands 13 and 14, contains a shorter β-strand (Ala610–Leu613) than predicted (Ile609–Leu614), whereas Thr692–Tyr697 in L8 forms a β-strand in the AlphaFold3 model but adopts a disordered loop conformation experimentally. Additional differences occur within the plug domain, where the experimental structure contains shorter β-strands spanning Arg247–Leu251 and Gly266– Val270 than predicted (Ile245–Arg252 and Gly266–Leu272, respectively). The Ala147–Asp149 and Ser256–Tyr259 segments similarly form short α-helical turns in the model but are disordered in the experimental structure (Fig. 4). Together, these data validate the global structural prediction while demonstrating that experimentally determined models are required to capture the conformational plasticity of extracellular loops and other membrane-proximal regions that may be critical for TBDT function.

**Figure 4.**
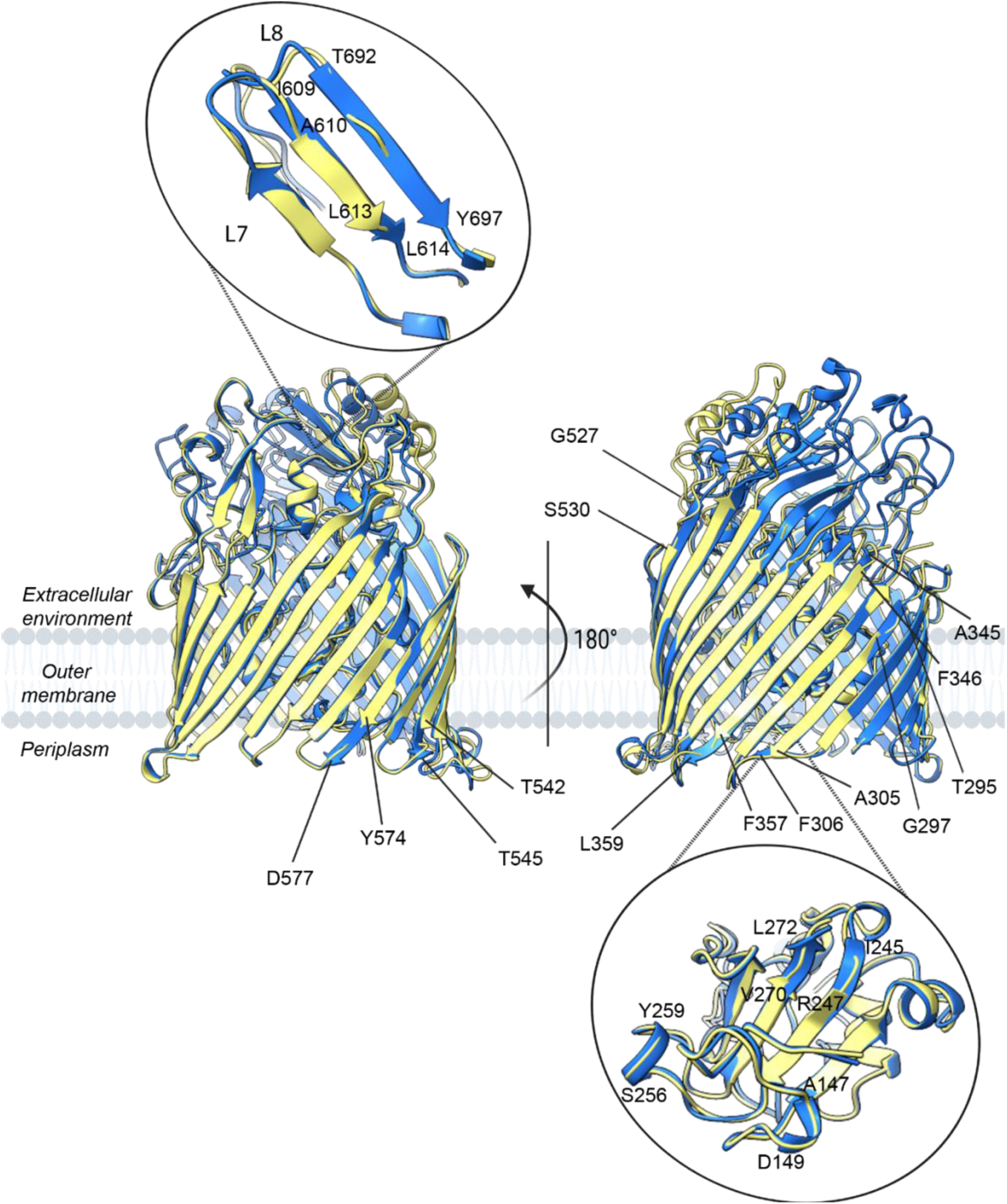
Structural comparison of the cryo-EM and AlphaFold 3 models of PA2590. Superposition of the cryo-EM structure (yellow) and AlphaFold 3 model (blue). The N-terminal region (residues 1-133) was omitted from the AlphaFold 3 model for clarity. The oval indicates extracellular loops 7 and 8, which are enlarged and rotated in the inset to highlight differences between the experimentally determined and predicted conformations. For detailed comparison, secondary-structure segments comprising residues 598-616 and 689-698 were superimposed separately, yielding an RMSD of 0.679 Å across 19 Cα atom pairs.

Similarity searches identified previously characterized TBDTs with diverse functions and limited sequence similarity to PA2590. The closest match was the catecholate siderophore/colicin receptor CirA from *Escherichia coli* (PDB 2HDI; Z-score, 38.5; RMSD, 2.3 Å over 559 aligned residues; 21% sequence identity), followed by the vitamin B12 transporter BtuB (PDB 1NQE; Z-score, 37.3; RMSD, 2.5 Å over 533 aligned residues; 21% sequence identity). Strong structural similarity was also observed with the *P. aeruginosa* catecholate siderophore receptors PirA (PDB 5FP2, chain B; Z-score, 34.6; RMSD, 2.3 Å over 509 aligned residues; 19% sequence identity) and PfeA (PA2688; PDB 5M9B; Z-score, 33.5; RMSD, 3.1 Å over 582 aligned residues; 18% sequence identity). Despite this overall fold conservation, PA2590 differs substantially from these TBDTs in the organization and length of its extracellular loops. In particular, in PA2590 the L7 (592–631) and L8 (662–697) form a prominent surface-exposed loop pseudodomain (LPD) surrounding the extracellular barrel entrance (Fig. 4). This organization differs markedly from the LPD architecture of PfeA and PirA, in which L5, L7, and L8 contribute to a single LPD. These structural differences argue against assigning the physiological ligand of PA2590 solely on the basis of global structural similarity and instead suggest that PA2590 may recognize a distinct substrate.

Among the 35 predicted TBDTs in *P. aeruginosa*, PA2590 shares close sequence similarity (64% ID) with the uncharacterized receptor PA2089. Not unexpectedly, the AF3 model of PA2089 also predicts two distinct LPDs formed by L4/L5 and L7/L8, matching the organization observed in PA2590. These four LPD-forming loops have identical lengths in the two proteins, whereas the cumulative absolute difference across all 11 extracellular loops was only two residues, arising from single-residue differences in L2 and L11 (Supplementary file PA2590LoopAnalysis.xlsx). This structural correspondence identifies PA2089 as a particularly intriguing candidate for future functional investigation and suggests that comparison of extracellular loop architectures may provide a useful basis for grouping uncharacterized TBDTs according to their potential substrate-recognition mechanisms.

### Structural and Functional Analysis of *P. aeruginosa* 12-Stranded OMBB proteins

Among the 105 proteins analysed, 17 were predicted to contain a single-polypeptide 12-stranded OMBB (Supplementary Table SX, sheet “Annotations of individual prot”). At the barrel-domain level, ECOD classified seven as OmpA-like domains (PA1763, PA1951, PA2421, PA2474, PA3772, PA4824 and PA5325), five as generic porin domains (PA1106, PA2064, PA4390, PA4391 and PA4579), three as autotransporter domains (PA0328, PA3535 and PA5112), and two as Tsx-like channels (PA0165 and PA0234) (Supplementary Table SX, sheet “Domain types and distribution”). Thus, a 12-stranded OMBB represents a structural class encompassing several functionally distinct protein families. Because domain-level classification may assign an autotransporter translocator domain simply as a porin-like barrel, we next examined the full-length architectures of the 12-stranded OMBB proteins for the combination of an N-terminal passenger region and a C-terminal translocator domain characteristic of classical autotransporters.

Type Va autotransporters, the best-characterized members of the type V secretion system, share a conserved architecture comprising an N-terminal passenger domain and a C-terminal translocator domain that forms a 12-stranded outer-membrane β-barrel^53,54^. Following translocation across the outer membrane, the passenger either remains associated with the cell surface or is released into the extracellular milieu, where it can mediate diverse functions, including adhesion, biofilm formation, host colonization, immune modulation and cytotoxicity^54^.

Our analysis identified six members of 12-stranded OMBB group in *P. aeruginosa* PAO1 (PA0328, PA3535, PA4390, PA4391, PA4579 and PA5112) that are predicted to adopt the canonical two-domain architecture of classical autotransporters. Three of these assignments are supported by previous experimental characterization. PA0328/AaaA is a cell-surface-tethered, arginine-specific aminopeptidase that promotes peptide utilization and virulence^47^. PA3535/EprS is a serine protease that activates protease-activated receptor signalling and induces inflammatory responses^55^; it has additionally been implicated in nutrient utilization, multiple virulence-associated phenotypes, and pathogenicity in a mouse infection model^12^. PA5112/EstA is a surface-displayed esterase required for normal rhamnolipid production, motility, and biofilm formation^56^. Its full-length crystal structure revealed a globular esterase passenger connected to a 12-stranded β-barrel, with the active site positioned on the apical surface of the passenger domain^57^. During this study, PA0328/AaaA was successfully expressed and purified at scale, and cryo-EM grids were prepared for single-particle analysis. However, publication of its structure, which confirmed the autotransporter architecture and elucidated the molecular basis of substrate recognition and catalysis^58^, led us to discontinue further structural characterization of this target. Consequently, PA3535/EprS remained the only experimentally characterized *P. aeruginosa* autotransporter in this group for which no structure was available.

The remaining three proteins, PA4390, PA4391, and PA4579, have not been experimentally characterized and may represent additional, functionally distinct members of the *P. aeruginosa* autotransporter repertoire. AF3 models of PA4390 and PA4391 each comprise a compact, β-rich SH3-like N-terminal domain (residues ∼22–116 and ∼20–133, respectively) followed by a C-terminal DUF481 porin-like barrel. Family-level bioinformatic analysis has classified DUF481 proteins as putative 12-stranded outer-membrane pore-forming proteins^59^. However, this classification does not establish an autotransporter function for PA4390 or PA4391, nor can ligand-binding or catalytic activities be assigned to their N-terminal domains based solely on structural similarity. PA4579 has a substantially larger predicted N-terminal region (residues ∼25– 386), with ECOD annotations indicating fold-level similarities to zincin-like, cysteine-proteinase-like, and type III secretion system domains. InterPro^60^ additionally annotates residues 119–292 as an Lnb N-terminal periplasmic domain. These assignments derive from ECOD and InterPro classification resources^61,62^ and do not establish proteolytic activity, cellular topology, or surface exposure experimentally.

The divergent predicted passenger-domain architectures of these proteins suggest substantial functional expansion within the *P. aeruginosa* autotransporter repertoire but also highlight the limitations of computational annotation alone. We therefore selected two complementary experimental approaches: structural characterization of the well-established serine protease EprS (PA3535) to define the molecular basis of its autotransporter architecture and proteolytic activity, and functional characterization of the previously uncharacterized PA4579 to test whether its predicted protease-like architecture corresponds to enzymatic activity.

### Structural Basis of EprS Autotransporter Function and Proteolytic Activity

PA3535/EprS autotransporter has been shown to activate protease-activated receptors (PAR-1, PAR-2, and PAR-4) on human epithelial cells through proteolytic cleavage of their tethered ligand regions by subtilisin-like serine protease passenger domain^55,63^. This results in NF-κB activation and IL-8 production through PAR-2 signaling^55^. The disruption of the corresponding gene in *P. aeruginosa* resulted in a pleiotropic phenotype that included a growth defect on nutrient-limited medium and attenuated virulence, thus confirming the important role of this autotransporter for *P. aeruginosa* pathogenesis^63^.

We recombinantly expressed and purified EprS and determined its structure using single-particle cryo-EM. This analysis yielded a 2.34 Å reconstruction of the N-terminal passenger domain spanning residues 30 to 673, with residues 635 to 660 unresolved, and a 3.23 Å reconstruction of the C-terminal OMBB domain comprising residues 633 to 995. Fitting these domain structures into the full-length EprS reconstruction defined their relative arrangement within the intact protein (Fig. 5).

**Figure 5.**
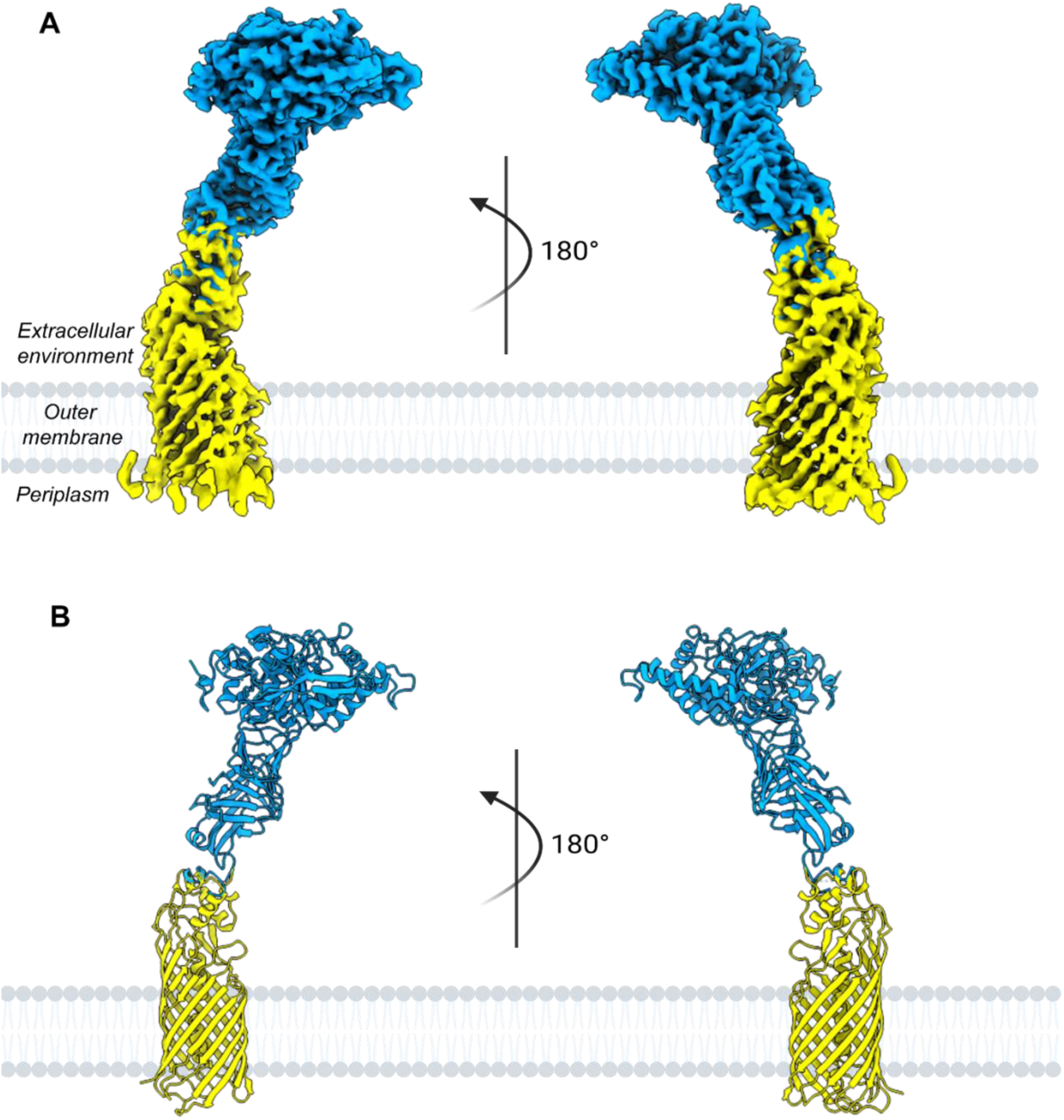
Cryo-EM structure of the mature EprS autotransporter (PA3535). (A) Composite cryo-EM density map of mature EprS shown in two orientations related by a 180° rotation. The passenger domain is coloured cyan, and the translocator domain, including the β-barrel and associated linker helices, is coloured yellow. (B) Cartoon representation of the composite atomic model shown in the same orientations and coloured as in A. The passenger-and translocator-domain reconstructions reached overall resolutions of 2.34 Å and 3.23 Å, respectively, and were combined to reconstruct the continuous architecture of mature EprS from the membrane-embedded β-barrel to the extracellular passenger domain. The passenger comprises an N-terminal globular protease domain followed by a β-helix stalk. The outer membrane is represented schematically in grey, and curved arrows indicate the 180° rotations.

The N-terminal passenger domain of EprS adopts a bipartite architecture consisting of a compact N-terminal globular peptidase subdomain and an elongated β-helix stalk (Fig. 5 and 6). The globular domain (residues 52–367) features an alternating arrangement of β-strands and α-helices connected by multiple flexible loops or short turns. This architecture is characteristic of the canonical subtilisin-like α/β fold, comprising a central twisted seven-stranded parallel β-sheet flanked by α-helices that form the proteolytic core. At the junction between the globular and stalk regions, a distinctive helix–turn–helix motif (residues 396 to 449) creates a finger-like projection that mediates the structural transition between the peptidase subdomain and the β-helical stalk. The latter subdomain (residues 454 to 571) features a right-handed parallel β-helical structure composed of repeating β-strand rungs and short connecting loops, followed by a 3₁₀-helix (residues 579 to 585) and a terminal β-hairpin motif (residues 613 to 619) that caps the distal region.

**Figure 6.**
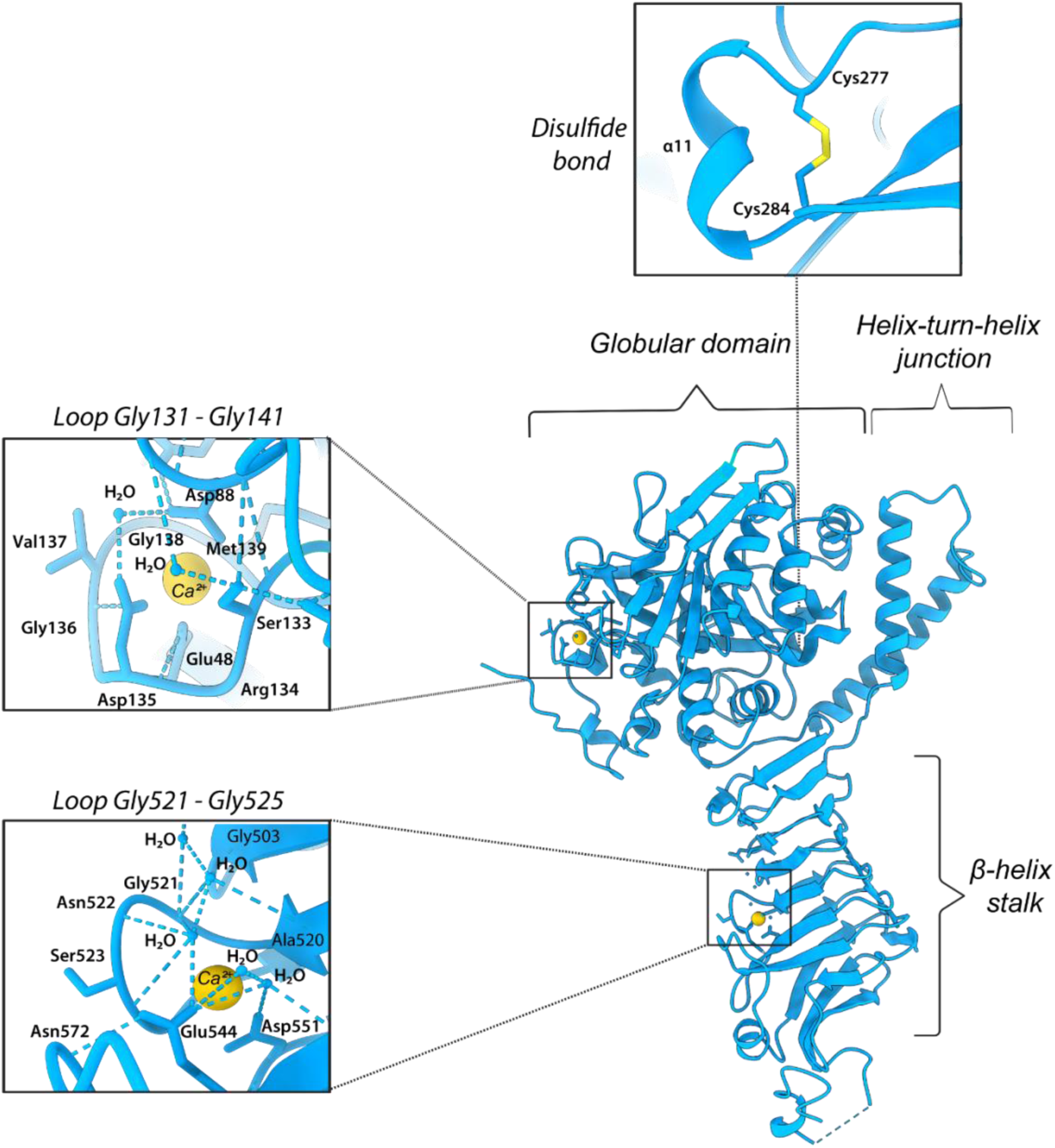
Structural features of the EprS passenger domain. Two additional densities in the cryo-EM map of the EprS passenger domain were modelled as bound Ca²⁺ ions based on their coordination environments. The passenger domain is shown in cartoon representation (light blue), with Ca²⁺ ions depicted as yellow spheres and the disulfide bond in gold. (Top inset) A conserved disulfide bond between Cys277 and Cys284 connects α11 to an adjacent β-strand, reinforcing the α/β interface of the globular domain. (Middle inset) Within the globular subdomain, a Ca²⁺ ion is coordinated by backbone and side-chain oxygen atoms from Glu48, Asp88, Ser133, Asp135, Val137, and Met139, stabilizing the Gly131–Gly141 loop in a distorted octahedral, six-coordinate geometry. (Bottom inset) A second Ca²⁺-binding site is located along the β-helix stalk near the Gly521–Gly525 loop. The ion adopts a six-coordinate geometry involving backbone carbonyl oxygen atoms from Ala520 and Ser523, carboxylate oxygen atoms from Glu544 and Asp551, and two ordered water molecules. Together, the two Ca²⁺-binding sites and disulfide linkage are positioned to stabilize local loop conformations and reinforce the passenger-domain architecture. Dashed lines indicate Ca²⁺-coordination and hydrogen-bonding contacts of ≤3.5 Å.

We identified additional density that we interpreted as ions that would stabilize flexible regions within the passenger domain (Fig. 6). One of the ions resides in the globular domain, where it forms ionic interactions with backbone carbonyl oxygens of residues Ser133, Val137, and Met139 belonging to a surface-exposed loop spanning residues 131 to 141, as well as with the side-chain oxygens of residues Asp135 and Glu48, completing a distorted octahedral, six-coordinate geometry with Asp88. A second ion is positioned along the β-helix stalk near Gly521– Gly525, where it is coordinated by main-chain carbonyl oxygens from Ala520 and Ser523, side-chain carboxyl groups from Glu544 and Asp551, and two ordered water molecules. These sites display coordination distances of approximately 2.1–2.6 Å, consistent with canonical six-coordinate, oxygen-ligated Ca²⁺ centers described in other subtilisin-like autotransporters^64^. Both ions act as structural stabilizers of loop flexibility rather than as catalytic cofactors that can reinforce the overall integrity and rigidity of the passenger domain architecture.

The structure of PA3535 also features a disulfide bond between Cys277 and Cys284 in the globular domain that anchors helix α11 to a neighbouring β-strand and likely stabilizes the local α/β interface (Fig. 6). The close spacing of these cysteines generates a compact disulfide-constrained segment compatible with autotransporter passenger translocation, whereas larger or more rigid disulfide-bonded loops can impede secretion^65^. An analogous disulfide bond between Cys288 and Cys295 is present in the subtilisin-like autotransporter Ssp from *Serratia marcescens*^64^, suggesting that this local stabilizing element may be shared among a subset of subtilisin-like autotransporters.

The C-terminal translocator domain of EprS forms a membrane-embedded OMBB spanning residues 633 to 995 (Fig. 7). A short helix bundle (residues 633 to 681) is positioned at the interface between the passenger domain and the outer membrane, which is followed by an elongated α-helix (residues 684 to 708) that extends into and occludes the barrel lumen. This helix runs axially inside the OMBB and occupies the center of the lumen, making close contacts with interior β-strands and blocking through-pore continuity. As predicted, the β-barrel itself comprises 12 antiparallel β-strands forming a single closed cylinder, with adjacent strands connected by alternating extracellular and periplasmic loops rather than short β-turns. The external set of loops is relatively long and flexible, consistent with the architecture of other autotransporter translocator domains^57,66^. The β-barrel exhibits an internal diameter of approximately 25 Å (radius ≈ 12.5 Å), typical of autotransporter pores capable of accommodating helical linker or folded passenger segments within the lumen^57,66^.

**Figure 7.**
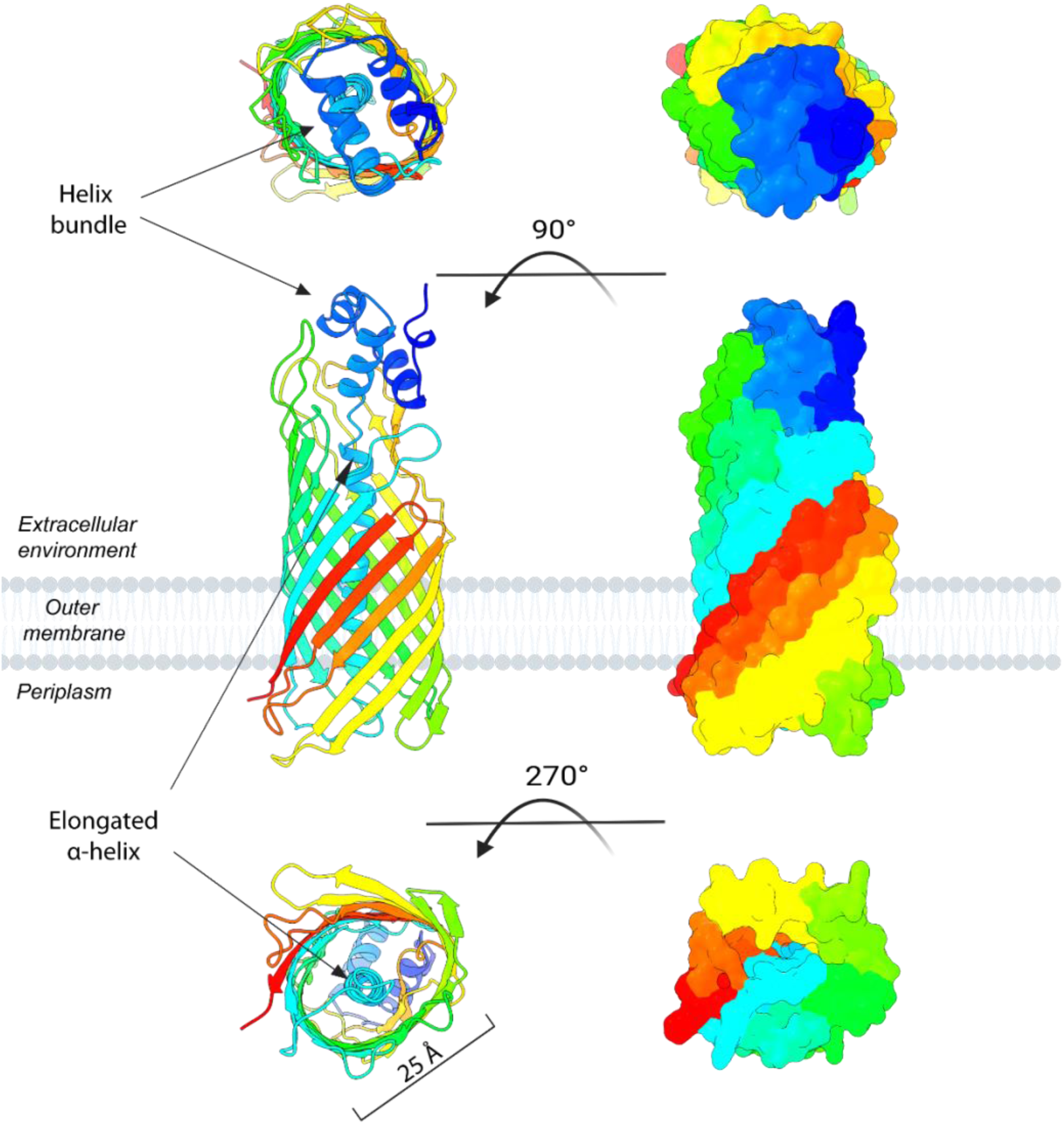
Architecture of the EprS translocator domain. The experimentally determined cryo-EM model of the C-terminal translocator domain (residues 633–995) is shown in cartoon (left) and molecular-surface (right) representations viewed from the extracellular side (top), parallel to the outer-membrane plane (middle), and from the periplasmic side (bottom). A short helix bundle (residues 633–681) lies at the interface between the passenger domain and the outer membrane and is followed by an extended α-helix (residues 684–708) that runs axially through and occludes the lumen of the 12-stranded antiparallel β-barrel. The barrel has an internal diameter of approximately 25 Å. The structure is coloured from the N-terminus (blue) to the C-terminus (red), and the outer membrane is represented schematically in grey.

The EprS’s protease domain belongs to the peptidase family S8 (MEROPS S08.162; InterPro IPR034061). Although subtilases and trypsin-like serine proteases adopt unrelated folds, both typically employ a Ser–His–Asp catalytic charge-relay system, representing a classic example of convergent evolution^67^. A Foldseek search^68^ identified the passenger domain of Ssp, a subtilase autotransporter from *Serratia marcescens* (PDB 8E7F)^64^, as a close structural match to the PA3535 passenger domain. Both passenger domains display a similar overall architecture, comprising an N-terminal subtilase α/β domain followed by a C-terminal three-stranded β-helical stalk. Superposition of the conserved regions yielded an RMSD of 0.90 Å across 387 Cα pairs after iterative outlier rejection (Fig. 8A, B). The alignment maps PA3535 Asp79, His122 and Ser308 onto the Ssp catalytic residues Asp76, His112 and Ser341, respectively, supporting their assignment as the EprS catalytic triad (Fig. 8B, C). Consistent with this assignment, substitution of EprS Ser308 with alanine was previously shown to markedly reduce proteolytic activity^55^. In addition, InterPro^60^ server identifiers Ser188–Trp189–Gly190–Asn237 residues adjacent to the catalytic cleft (Fig. 8D) as nucleophilic-elbow and oxyanion-hole motifs characteristic of subtilisin-like proteases and other α/β-hydrolase enzymes, which stabilize the tetrahedral intermediate and orient the substrate during catalysis^67,69^. The backbone carbonyl of Ser188 and the side-chain hydroxyl of Ser308 form two reciprocal polar contacts (Ser308 OG–Ser188 O and Ser188 OG–Ser308 O, 3.1 and 2.6 Å, respectively), while all other catalytic distances exceed typical hydrogen-bonding ranges, probably due to the apo state of EprS structure. Only the Asp79 OD2–His122 ND1 pair remains within bonding distance (2.7 Å), consistent with partial retention of the charge-relay alignment (Fig. 8D). Together, these measurements indicate a relaxed, pre-reactive geometry in which the catalytic triad is largely disengaged, while the nucleophilic-elbow loop remains pre-organized to stabilize the transition state upon substrate binding.

**Figure 8.**
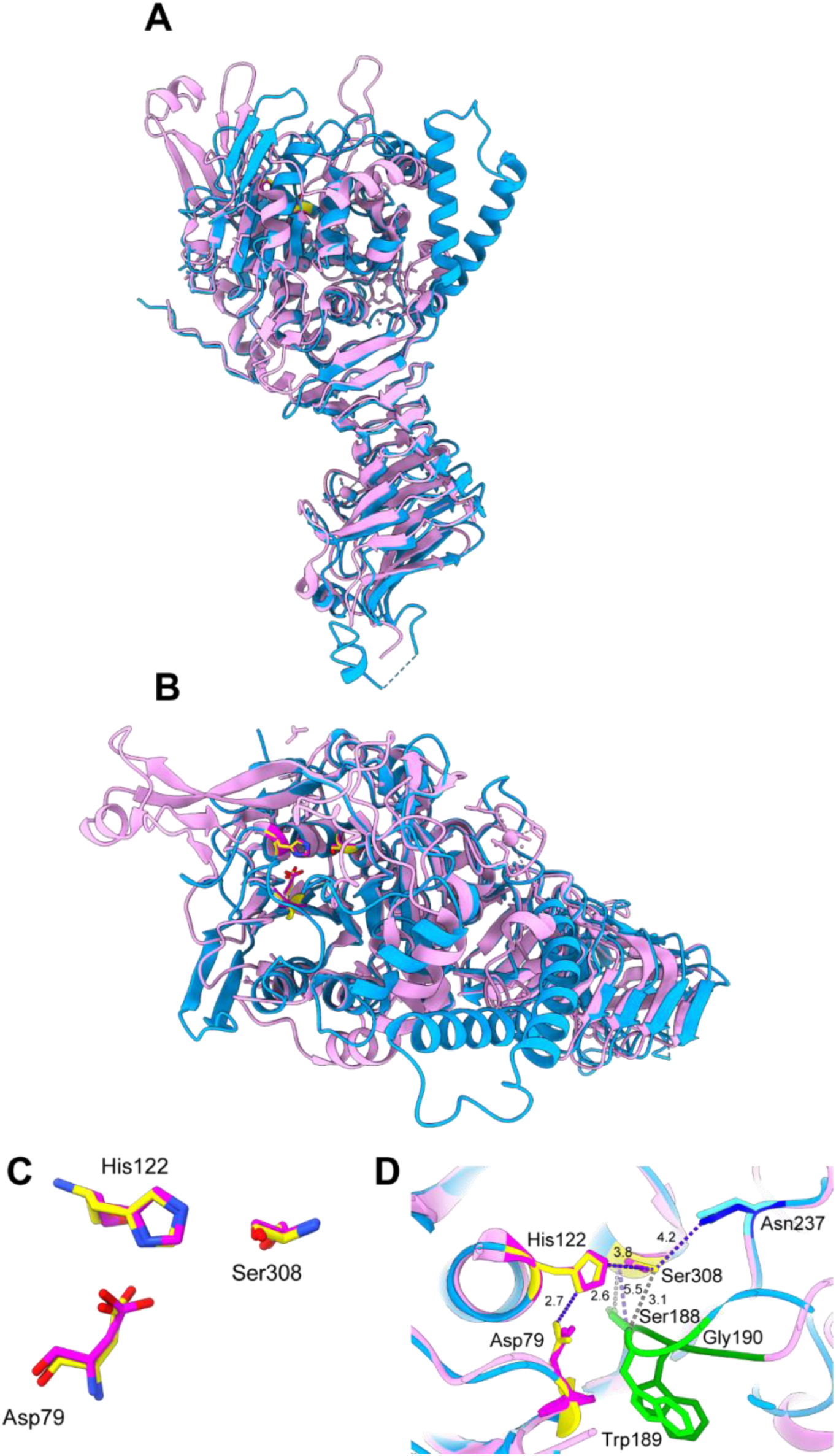
Structural alignment of EprS (PA3535) and Ssp (PDB ID: 8E7F). (A) Side view of the backbone superposition of the EprS passenger domain (sky blue) and Ssp (pink; PDB ID: 8E7F). (B) View from the side highlighting the globular catalytic domain in the same superposition. (C) Close-up of the catalytic triad region extracted from the aligned passenger domains. Residue numbering corresponds to EprS. (D) Detailed view of the putative active site in the superimposed structures. The catalytic triad residues are shown in yellow for EprS and magenta for Ssp; the nucleophilic-elbow loop (Ser–Trp–Gly) is colored lime in EprS and forest green in Ssp; and the Asn residue forming the oxyanion-hole stabilizer is shown in dark blue for EprS and cyan for Ssp. Observed hydrogen bonds are depicted as gray dashed lines, whereas nonbonded charge-relay distances are shown in purple. Residue numbering corresponds to EprS.

Next, we compared the EprS cryo-EM structure with the AF3 predicted model. Overall, the predicted model of EprS is in good agreement with the determined cryo-EM structure (Fig. 9A-D; Fig. 10A-B), manifesting in RMSDs of 0.53 Å for the passenger domain (residues 52–619) and 0.76 Å for the translocator β-barrel and plug region (residues 633–955). However, the RMSD without exclusion of atoms increased to 1.1 Å in the translocator domain, reflecting conformational flexibility in certain elements (Fig. 9C). Furthermore, several regions in EprS show local differences between the experimentally defined structure and the AF3 model, with shifts in the boundaries of secondary-structure elements (Fig. 10A). In the translocator β-barrel, the cryo-EM density resolves flexible surface loops in locations where AF3 predicts extended β-strands or more ordered elements. Specifically, the loops formed by residues Ala960 to Leu972 and Arg798 to Leu805 in the structure were predicted to be shorter and to encompass only residues 963 to 968 and 800 to 803, respectively. Also, a disordered region between residues Gly907 and Asp909 was predicted to form a short β-strand. Similarly, the loop between residues 820 and 833 was predicted to encompass the residues 824 to 827 (Fig. 10B). Observed differences are in line with our general observation mentioned above that the AF model overestimates β-strand continuity in solvent-exposed regions that are likely destabilized in the membrane environment. Together, these analyses indicate that the AF prediction accurately recapitulates the global fold of EprS (PA3535), while over-predicting the rigidity of the surface-exposed and membrane-embedded boundaries, where the cryo-EM data reveal greater conformational plasticity.

**Figure 9.**
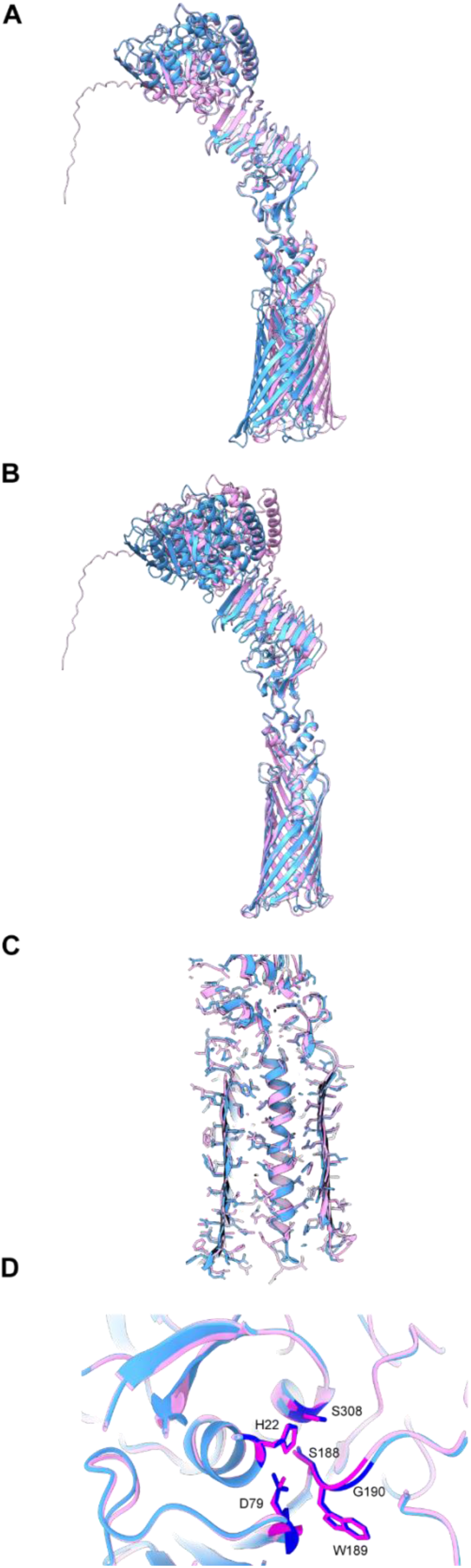
Structural comparison between the cryo-EM and AlphaFold3 models of PA3535. (A) Alignment of the passenger domains from the cryo-EM structure (blue) and AlphaFold3 prediction (pink) shows close backbone correspondence across the subtilisin-like core. (B) The β-barrel translocator domains, shown in the same coloring scheme, also superimpose with high overall agreement in the backbone architecture. (C) Close-up of the translocator and lumenal plug region using the superposition in B. Front-facing barrel strands are omitted from the display to expose the barrel interior, and side chains are shown as sticks (D) Close-up of the catalytic region using the passenger-domain superposition in A. Catalytic residues are shown as sticks, coloured bright blue in the cryo-EM-derived model and magenta in the AlphaFold3 prediction.

**Figure 10.**
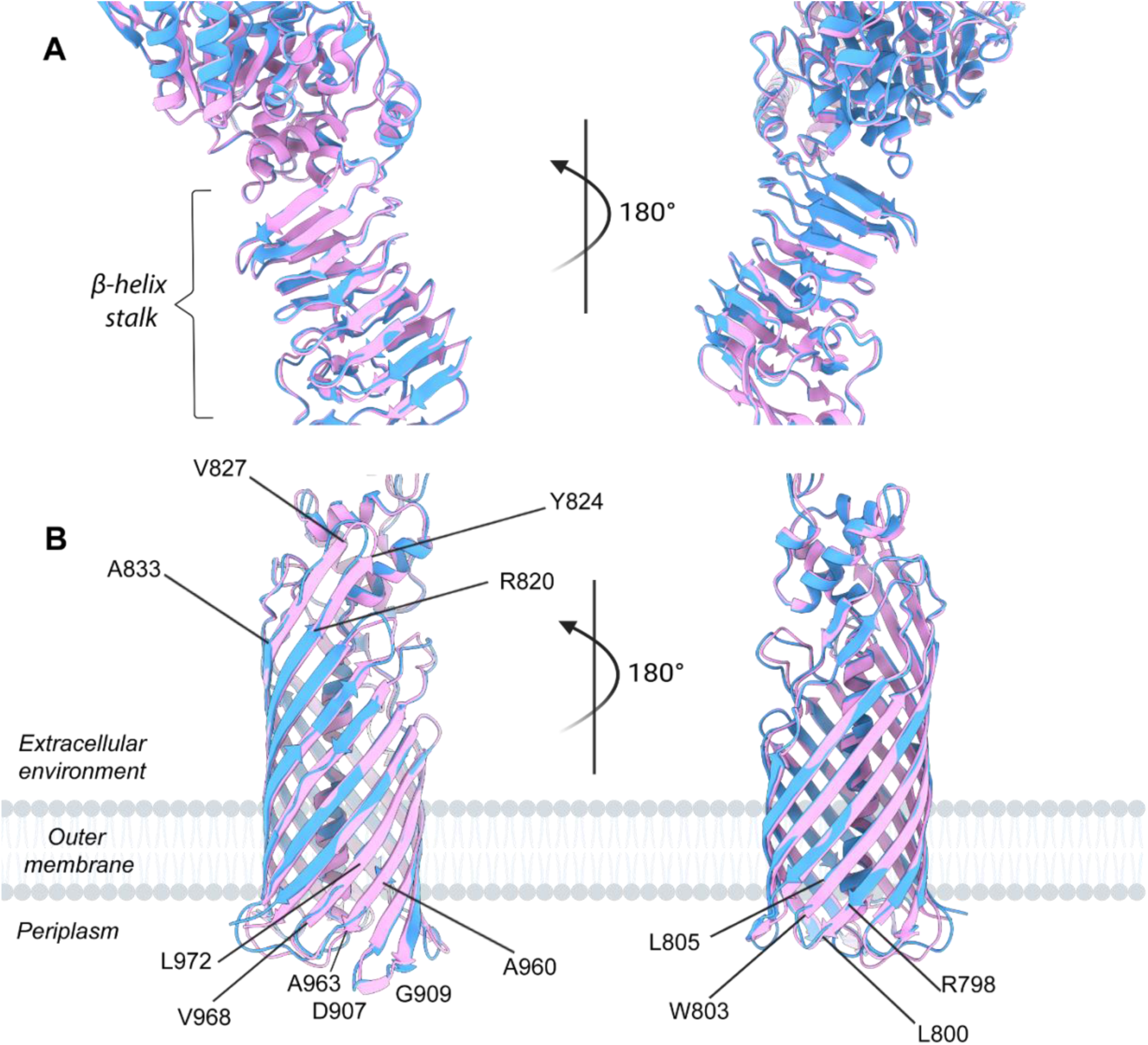
Secondary-structure deviations between the cryo-EM structure of PA3535 and the AlphaFold3 prediction. (A) Enlarged view of the aligned β-helical stalk from the cryo-EM model (blue) and the AlphaFold3 prediction (pink). (B) Alignment of the β-barrel translocator domains, shown in the same color scheme.

### PA4579 autotransporter exhibits phospholipase A1 activity *in vitro*

We next investigated PA4579, one of three previously uncharacterized PAO1 proteins predicted to adopt the general architecture typical of autotransporters. The AF3 model predicts a large, multidomain N-terminal region (residues 25–386) preceding the 12 strand OMBB (residues 387–618) (Supplementary Fig. 2). By analogy with characterized type Va autotransporters, we refer to these regions hereafter as the putative passenger and translocator domains, respectively.

The putative passenger domain was annotated only as a member of the DUF4105 family (Supplementary Fig. 2). Structural comparison of AF3 models, however, revealed close similarity between the PA4579 DUF4105 domain and the periplasmic domain of Lnb, a lipoprotein *N*-acyltransferase from *Bacteroides thetaiotaomicron*^70^. The predicted fold superimposed with an RMSD of approximately 1.7 Å over 114 Cα pairs following distance-based pruning, despite only 28% sequence identity (Supplementary Fig. 3). This comparison identified PA4579 residues Asn173, His150, and Cys256 as structural equivalents of the proposed Lnb catalytic triad Asn66– His49–Cys148^70^. Cys148 has been shown to be required for Lnb activity, whereas the roles of Asn66 and His49 remain inferred from structural modelling and sequence conservation^70^. These observations suggested that the PA4579 passenger domain might possess related phospholipid-modifying activity.

To test this hypothesis, we incubated purified wild-type PA4579 or the variant carrying C256A substitution with 1-palmitoyl-2-oleoylphosphatidylethanolamine (POPE) or 1,2-dioleoyl-sn-glycero-3-phosphoethanolamine (DOPE) phospholipid substrate in the presence of FSL-biotin as a candidate acyl acceptor and analyzed the reaction products by HPTLC (See Methods). A prominent product that co-migrated with the lysophosphatidylethanolamine (LPE) standard was detected in the reaction containing wild-type PA4579 and DOPE, whereas the corresponding band was markedly weaker or undetectable in case of POPE. The prominence of this product was significantly lower in case of C256A variant (Fig. 11). These results are consistent with Cys256-dependent deacylation of DOPE by PA4579 under the conditions tested. DOPE appeared as a more preferable substrate for PA4579 compared to POPE. This observation prompted us to suggest that PA4579 exhibits substrate selectivity toward PE species containing a monounsaturated oleoyl chain at the sn-1 position rather than saturated palmytoyl. Since we did not detect any product consistent with transfer of the released acyl group to FSL-biotin we concluded that PA4579 does not exhibit the acyl-transfer activity towards this candidate acceptor under used conditions.

**Figure 11.**
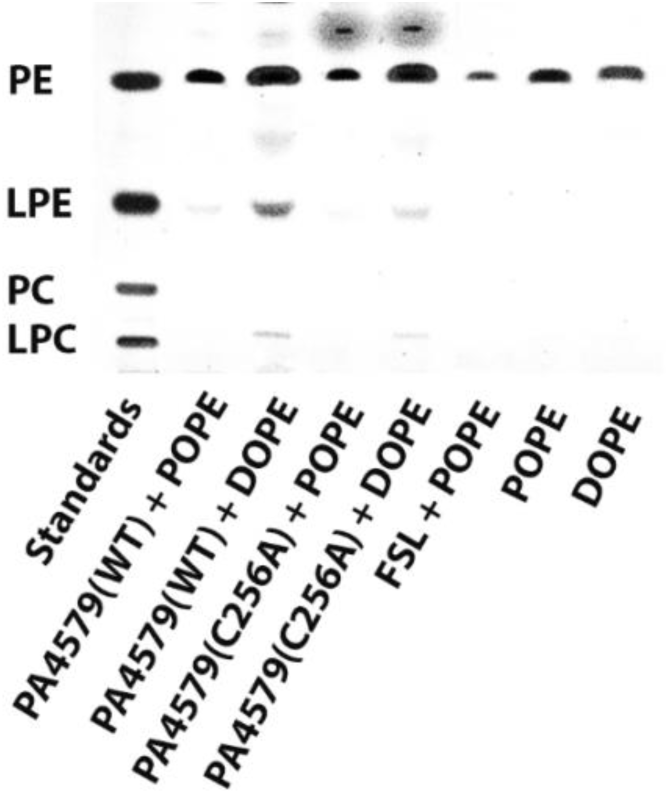
PA4579 catalyzes Cys256-dependent deacylation of DOPE. HPTLC analysis of POPE- and DOPE-containing samples, obtained from wild-type (WT) PA4579 or PA4579 C256A catalyzed reactions, using CuSO4-based staining. A no-enzyme reaction containing FSL-biotin and POPE (FSL + POPE), together with POPE- and DOPE-only controls, was included. PE, LPE, PC, and LPC standards were run simultaneously on the same HPTLC plate and are shown on the left. A prominent band corresponding to the LPE standard (same Retention factor (Rf)) was detected following incubation of DOPE with wild-type PA4579. The corresponding signal was substantially weaker with PA4579 C256A and negligible in the remaining reactions, supporting Cys256-dependent deacylation of DOPE by PA4579. HPTLC plates were visualized using excitation/emission wavelengths of 540/590 nm; a representative plate is shown here.

It is well established that fluorescent lipid analogs enable more sensitive detection of reaction products than the routine HPTLC analysis of unlabeled phospholipids. Therefore, to further examine PA4579 activity, we performed two independent assays using a defined fluorescent substrate, NBD-labelled PE [1-oleoyl-2-(12-NBD-dodecanoyl)-PE]. Wild-type PA4579 generated a strong fluorescent signal corresponding to a product migrating at the position corresponding to NBD-LPE, whereas no corresponding product was detected in the substrate-only control. Although a very low-level background signal was observed in the FSL-biotin plus NBD-PE control, the markedly stronger signal detected in the presence of wild-type PA4579 suggests that product formation was strongly dependent on the PA4579 catalysis and was unlikely to be attributable solely to background or non-enzymatic activities (Fig. 12A). Product formation was substantially reduced by the C256A substitution (Fig. 12A) and was not detected once double (C256A/H150A) and triple (C256A/H150A/N173A) substitutions were introduced (Fig. 12B). Because the NBD-labelled acyl chain occupies the *sn*-2 position of the substrate, its retention in the fluorescent lysophospholipid product NBD-PE is consistent with cleavage of the non-fluorescent *sn*-1 acyl chain. Based on these data we postulated that PA4579 exhibits phospholipase A1-like activity *in vitro*. The reduced activity of the C256 along with His150 or Asn173 substitutions suggest their role in this activity however the full assessment of the catalytic mechanism of PA4579 remains to be further characterised. Together, these structural and biochemical analyses identify PA4579 as a previously uncharacterized autotransporter-like protein with phospholipase A1-like activity *in vitro*. This finding provides experimental functional evidence for the DUF4105-containing passenger region of PA4579 and expands the known enzymatic diversity of the predicted *P. aeruginosa* autotransporter repertoire. Product formation in the absence of FSL-biotin suggests that this compound is not required for phospholipid deacylation and favours hydrolysis over completion of an Lnb-like acyl-transfer reaction to FSL-biotin under tested conditions. Nevertheless, acyl transfer to a different physiological acceptor cannot be excluded. Whether PA4579 is exposed at the bacterial surface and which phospholipids or potential acyl acceptors represent its physiological substrates remain important questions for future investigation.

**Figure 12.**
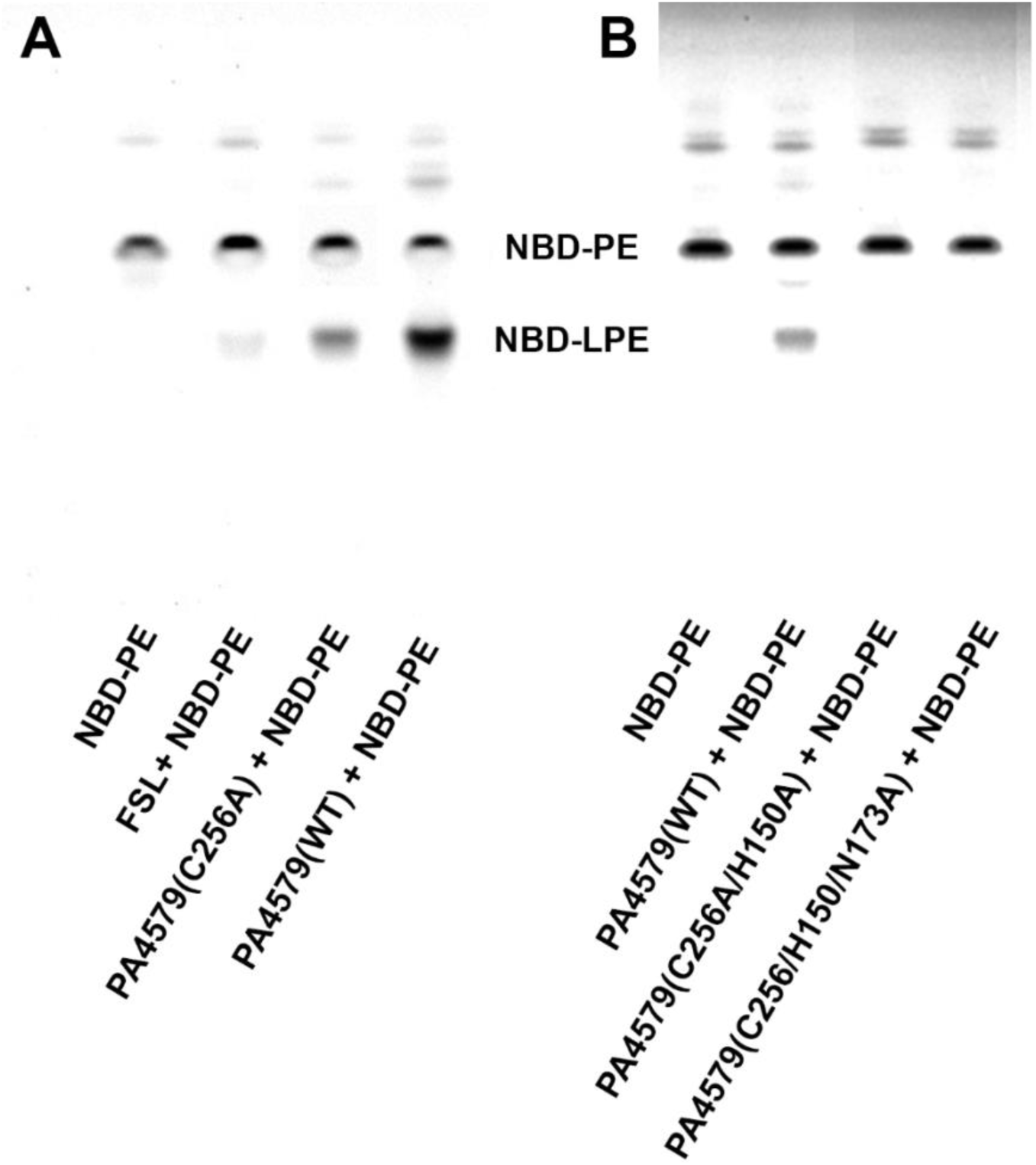
PA4579 generates NBD-LPE production from NBD-PE in the absence of an exogenous acyl acceptor. Fluorescence-detected HPTLC analysis of 18:1–12:0 NBD-PE incubated under the following conditions:(A) NBD-PE alone or in the presence of FSL-biotin, PA4579 C256A, or wild-type (WT) PA4579; (B) NBD-PE alone or incubated with wild-type PA4579, PA4579 C256A/H150A, or PA4579 C256A/H150A/N173A. An NBD-PE product corresponding to NBD-LPE was most prominent following NBD-PE incubation with wild-type PA4579. Product formation was markedly reduced with PA4579 C256A and was negligible in the FSL-Biotin + NBD-LPE control, while no NBD-LPE was detected in the combined double and triple variants containing C256A. These results support a central role for the Cys256 residue in PA4579-mediated phosphatidylethanolamine deacylation. HPTLC plates were visualized at excitation/emission wavelengths of 465/536nm; two representative plates are shown.

## Discussion

In this study, we combined sequence-and structure-based approaches to define the OMBB proteome of *Pseudomonas aeruginosa* PAO1. Our analysis demonstrates that proteome-scale structure prediction can substantially extend protein annotation beyond the limits defined by sequence similarity alone. We focused on proteins predicted to adopt closed, membrane-spanning antiparallel β-meander barrels, which is the canonical architecture of integral outer-membrane proteins in Gram-negative bacteria, and identified 105 proteins, each containing a single OMBB domain. These proteins could be further classified according to barrel size, β-strand organization, extracellular loop architecture, and accessory domain composition. By integrating complementary sequence-, HMM-, and predicted model-based approaches, we obtained a substantially more comprehensive and functionally informative view of the *P. aeruginosa* OMBB repertoire than is captured by conventional database annotations. Predicted model-based analysis uncovered remote relationships that were not readily detectable by sequence similarity, helped reconcile apparent discrepancies among OMBB predictions, and revealed that β-strand number, although diagnostic for several well-defined protein families, is insufficient on its own to establish functional identity. Instead, reliable functional classification often requires consideration of the complete structural architecture, including accessory domains and extracellular loop organization. The resulting structure-informed catalogue exposes functional diversity that is obscured by broad or incomplete database annotations and provides a generalizable framework for systematically reannotating bacterial OMBB proteomes. More importantly, it establishes a rational basis for prioritizing poorly characterized OMBBs for experimental structural and functional analysis, thereby transforming predicted structures from static annotations into testable hypotheses about protein function, substrate recognition, transport, and host–pathogen interactions.

Development of AI-driven protein structure prediction algorithms, such as AlphaFold, followed by several others (e.g. ESM-fold), has transformed protein structure prediction by providing reliable protein models, including for targets lacking close structural templates^37,40^. In the present study, analysis of predicted structures increased the number of OMBB proteins identified beyond those detected by BLASTP or HHpred and expanded the number of ECOD-classified domains from 159 based on sequence similarity to 201 based on structural comparison.

These findings demonstrate the value of AI structural model prediction for proteome-scale annotation, particularly for recognizing overall folds and detecting remote structural relationships that are not apparent from sequence alone. However, comparison with experimentally determined structures also showed that predicted models provide an incomplete and sometimes distorted representation of local architecture. Cryo-EM resolved flexible extracellular loops, β-strand boundaries, disordered regions, bound ions, and functionally relevant structural features were regularized, overextended, or absent in the AF3 models. Thus, structural prediction is a powerful tool for assigning global folds and generating experimentally testable hypotheses, whereas high-resolution structural studies remain essential for defining conformational heterogeneity, local intramolecular interactions, and the molecular features underlying substrate recognition and function.

Our analysis was deliberately restricted to integral outer-membrane β-barrels (OMBBs) in which a single polypeptide chain forms a complete membrane-spanning β-barrel. This definition provides a clear structural criterion for proteome-wide identification but necessarily excludes OMBB assemblies in which the transmembrane barrel is formed through oligomerization of multiple polypeptide chains. For example, *P. aeruginosa* OprM (PA0427) forms a trimeric outer-membrane channel in which each protomer contributes four β-strands to a shared 12-stranded transmembrane barrel^71^. Such oligomeric architectures cannot be reliably identified by evaluating individual polypeptides for barrel closure and instead require explicit consideration of quaternary structure and subunit assembly. A second class of outer-membrane proteins not captured by our analysis comprises OM-anchored lipoproteins, exemplified by *P. aeruginosa* LolB (PA4668), which functions in the terminal stage of outer-membrane lipoprotein localization^72^. Unlike integral OMBBs, LolB is anchored to the membrane through an N-terminal lipid moiety and does not form a complete membrane-spanning β-barrel. Structurally characterized LolB homologues instead adopt an incomplete β-barrel-like fold that creates a hydrophobic cavity capped by an α-helical lid^73,74^. Thus, although both oligomeric OMBB assemblies and OM-anchored lipoproteins constitute important components of the *P. aeruginosa* OM proteome, their distinct topological and structural principles require analytical approaches beyond the single-chain barrel criterion used here. These exclusions define the scope of the present catalogue while also highlighting important classes of OM proteins that remain to be systematically integrated into future structural model-guided proteome-wide analyses.

To assess the power of using structural models, we compared the list of predicted OMBB containing proteins to the prediction provided by IsItABarrel program and database (version from September 2026)^75^, originally built without using AF3 models. Assessing the proteome of *P. aeruginosa* PAO1 by IsItABarrel identified 130 proteins containing OMBB domain. In the case of 101 (80%) proteins, this prediction coincided with our analysis. However, for most of the remaining 29 proteins, the IsItABarrel server’s prediction contradicted our ECOD-based analysis of their 3D models built with AF3. The structures of nine proteins in this group have been determined experimentally, and another eighteen proteins have statistically significant matches to structures in the PDB detected using BLAST. The analysis of these proteins revealed that the structure of only one of these proteins -PA0423 (PDB ID: 7BWL), contains an eight-stranded, antiparallel β-barrel domain which, however, belongs to the lipocalin family^76^, which is not an OMBB in ECOD classification. Another two proteins predicted by IsItABarrel and not identified by our analysis are homologous to experimentally characterized β-barrel secretin domains forming multimeric β-barrels. As mentioned earlier, multimeric β-barrels were intentionally excluded from our study. In two cases where experimentally characterized structure of homologous protein was not detected by BLAST, predicted structural models with high confidence scores did not contain β-barrel domains; however, further experimental analysis is required to characterize the molecular structure of these proteins. Detailed information on predictions from IsItABarrel that are not congruent with our predictions is provided in Supplemental Table SX.

Beyond defining a list of *P. aeruginosa* proteins with OMBB topology, our analysis revealed that the major OMBB classes are associated with characteristic accessory-domain architectures. Specifically, all 35 predicted 22-stranded TonB-dependent transporters (TBDT) contained a lumen-occluding plug domain, whereas proteins with predicted 16-stranded OMBB domains of the Omp85 superfamily also featured periplasmic POTRA domains. Within the group with predicted 12-stranded OMBB, autotransporters formed a distinct architectural subgroup in which a C-terminal translocator barrel was linked to an N-terminal passenger domain. Additional modules, including N0 and prealbumin-like domains, further expanded the range of class-specific domain combinations. Together, these recurring combinations indicate that diversification of the *P. aeruginosa* OMBB proteome has occurred not only through variation in OMBB size, but also through the modular association of conserved membrane-embedded scaffolds with structurally distinct accessory domains. Structural model-based comparison was particularly valuable for resolving these relationships when sequence conservation alone was insufficient for confident annotation.

Our structural analysis of *P. aeruginosa* proteins predicted to contain OMBBs highlights both the power and limitations of fold-based functional assignment. Structural characterisation of PA2590 confirmed the predicted 22-stranded OMBB, capped by a plug domain and associated with an N-terminal TonB-related region, establishing the canonical architecture of a TonB-dependent transporter and providing structural support for reclassifying PA2590 from a conserved hypothetical protein. However, these structural features alone do not permit confident assignment of its physiological ligand. In TBDTs, substrate specificity is determined largely by the sequence and architecture of the extracellular loops and by features of the exposed plug-domain surface. These regions also showed the greatest divergence between the experimentally determined and predicted structures. In the ferric-enterobactin receptor PfeA, for example, the extracellular loops form an initial ligand-recognition site before the siderophore is repositioned into a deeper binding site adjacent to the plug domain^77^. Comparison of extracellular loop-length profiles across the PAO1 TBDT repertoire identified PA2089, which shares 65% sequence identity with PA2590, as its closest paralogue, whereas PfeA and PirA were the closest receptors with experimentally established catecholate-siderophore specificities. Notably, differences in loop-pseudodomain (LPD) organisation argue against inferring ligand specificity from overall structural similarity alone. These relationships may instead reflect shared structural constraints imposed by ligand dimensions or mechanisms of initial substrate capture at the cell surface. The distinctive loop architecture of PA2590 therefore provides a rational basis for prioritising bulky ferric catecholate xenosiderophores as candidate ligands, while leaving open the possibility that this transporter recognises a different, as-yet-unidentified substrate. This distinction is particularly relevant because PA2590 was recently identified as a virulence determinant in a murine model of ocular infection, although its proposed roles in iron or cobalamin acquisition remain unconfirmed^52^. Direct ligand-binding and transport studies will therefore be required to connect the structural features of PA2590 to its physiological function and to determine whether its distinctive extracellular loop architecture encodes a previously unrecognised substrate specificity. Importantly, our analysis also identified LPD-like structural modules in several other TBDTs and in some additional predicted OMBBs, suggesting that this architectural feature may represent a broader determinant of ligand recognition across the *P. aeruginosa* OM proteome. The recurrence of these modules, together with their pronounced structural variation, highlights the potential value of LPD architecture as an additional layer of structure-based functional annotation beyond global fold classification. Systematic identification and comparative analysis of LPD-like modules may therefore provide a powerful approach for predicting substrate classes and prioritising experimentally tractable candidates within the expanding OMBB repertoire. Further investigation of these modules and their contributions to ligand recognition is currently underway.

The analysis of *P. aeruginosa* proteins predicted to contain 12-stranded OMBBs reveals a distinct strategy of structural and functional diversification. Classical type Va autotransporters comprise an N-terminal extracellular passenger domain linked to a C-terminal 12-stranded translocator OMBB, with an α-helical segment occupying the barrel lumen during or following passenger-domain secretion^54^. Our analysis identified at least three previously uncharacterized proteins with this predicted autotransporter architecture, distinguished by markedly different passenger-domain structures, suggesting substantial functional diversification within this class.

In *P. aeruginosa* EprS (PA3535), the prediction model correctly captured the translocator OMBB and the overall organisation of the passenger domain into a subtilisin-like protease domain followed by an extended β-helical stalk. The cryo-EM structure of EprS, however, revealed structural features that could not be defined confidently from prediction alone, including the organisation of flexible surface loops, structural Ca²⁺-binding sites, an intramolecular disulfide bond stabilising the protease domain, and the geometry of the catalytic centre. These observations connect the previously established functions of EprS in protease-activated receptor cleavage, inflammatory signalling, and virulence to a defined molecular architecture^55,63^. Thus, although structure prediction can accurately identify the overall architectural class and guide construct design, experimental structure determination remains essential for resolving catalytic geometry, metal coordination, covalent stabilisation, and conformational features relevant to protein secretion and activity.

Functional analysis of the previously uncharacterized PA4579 further illustrates how examination of full-length protein architecture can reveal unexpected biochemical relationships that are not apparent from barrel-domain classification alone. Although ECOD classified its C-terminal 12-stranded OMBB as a generic porin domain, the presence of a substantially larger N-terminal region suggested an autotransporter-like architecture. This region, previously annotated as DUF4105, showed structural similarity to Lnb, a lipoprotein *N*-acyltransferase from *Bacteroides*, and contained a corresponding set of candidate catalytic residues. Lnb catalyses a reaction analogous to that of the canonical bacterial lipoprotein *N*-acyltransferase Lnt, despite being evolutionarily and structurally distinct^70^. Loss of NBD-phosphatidylethanolamine conversion into NBD-lysophosphatidylethanolamine by the PA4579 Cys256Ala variant and by combination variants containing this substitution supports a central role for Cys256, consistent with its predicted role within the catalytic site. However, the formation of NBD-lysophosphatidylethanolamine in the absence of detectable acyl transfer to FSL-biotin suggests that, under conditions tested, PA4579 primarily promotes phosphatidylethanolamine hydrolysis (acyl chain cleavage at sn-1 position) rather than transacylation reaction and, therefore, does not establish completion of an acyl-transfer reaction. Additionally, the preferential deacylation of DOPE over POPE *in vitro* suggests that PA4579 may exhibit substrate specificity influenced by the acyl chain composition of phosphatidylethanolamine at sn-1 position, although the basis and physiological relevance of that preference remain to be established. Given the asymmetric organization of the outer membrane, in which phosphatidylethanolamine is enriched in the periplasm-facing inner leaflet whereas the outer leaflet is dominated by lipopolysaccharides, we predict that orientation of PA4579 places its catalytic site in a position compatible with access to PE in the inner leaflet. This topology therefore provides a plausible structural basis for PA4579-mediated PE deacylation *in vivo* and is consistent with its phospholipase A1-like activity observed *in vitro*. We therefore interpret the observed activity as phospholipid deacylase activity consistent with phospholipase A1-like cleavage, which could represent the first step of an acyl-transfer reaction. More broadly, these findings illustrate an important principle of structure-guided functional annotation: structural similarity can reveal a conserved catalytic scaffold and nominate residues for experimental interrogation but does not establish conservation of the complete biochemical reaction. Together, the EprS and PA4579 analyses demonstrate the complementary strengths of computational prediction and experimental validation. Predicted structures provide a framework for architectural classification and hypothesis generation, whereas experimental structural and biochemical analyses are required to define functional boundaries, catalytic mechanisms, and physiological relevance.

To conclude, our findings support a model in which OMBB evolution preserves a structurally constrained membrane-embedded scaffold while diversifying the regions that determine molecular function. The OMBB provides a robust framework for membrane insertion and persistence within the asymmetric outer membrane, whereas extracellular loops, lumenal plugs, accessory domains, and autotransporter passenger regions provide structurally adaptable surfaces for substrate capture, energy coupling, secretion, signalling, and catalysis. This organisation helps explain both the strengths and limitations of current structural model-prediction approaches: global OMBB architecture can often be predicted with high confidence, whereas the flexible and functionally specialised regions that encode molecular specificity remain more difficult to resolve and interpret. Our results therefore support a hierarchical approach to OMBB annotation, integrating sequence similarity for close relationships, profile-based methods for remote homology, predicted structural model analysis for topology and domain organisation, and targeted structural and biochemical experiments for functional validation.

Applied at the proteome scale, this strategy substantially expands the recognised OMBB repertoire of *P. aeruginosa* and transforms a largely sequence-defined collection of proteins into a structure-informed catalogue of membrane architectures and functional hypotheses. The identification of previously unrecognised domain organisations, divergent extracellular loop architectures, and catalytic passenger domains demonstrates that substantial functional information remains hidden within poorly annotated bacterial outer-membrane proteomes. Importantly, these features provide experimentally tractable entry points for investigating nutrient acquisition, secretion, virulence, host interactions, and antimicrobial susceptibility. By coupling scalable structure prediction with focused experimental validation, our study establishes a general framework for converting proteome-scale structural models into mechanistically testable hypotheses and provides a roadmap for systematically expanding functional annotation of the bacterial outer membrane. This approach should be broadly applicable beyond *P. aeruginosa*, particularly to pathogens whose outer-membrane proteomes remain dominated by proteins of unknown or incompletely defined function.

## Methods

### Identification of *P. aeruginosa* OMBBs and their domain architectures based on ECOD hierarchy

We used a combination of sequence similarity and automated structure analysis of AI-generated 3D models of *P. aeruginosa* proteins to identify all outer membrane meander beta barrels (OMBBs) in its proteome.

First, protein sequences from *P. aeruginosa* reference proteome (strain ATCC 15692 / DSM 22644 / CIP 104116 / JCM 14847 / LMG 12228 / 1C / PRS 101 / PAO1, ID: UP000002438, version from 2025/06/30) were downloaded from the UniProt database. These sequences were compared to the sequences of entries from the ECOD database using BLASTP^48^ and HHPred^43^ (sequence-and HMM-based algorithms, respectively). The proteins with significant (E-values <= 0.001) sequence similarity to any ECOD entry from Outer membrane meander beta-barrels X “possible homology” level were collected.

In the next step, structural models of proteins from the same proteome were generated using AlphaFold 3, either through a local installation on the University of California, Riverside High-Performance Computing Cluster (UCR HPCC) or via the online AlphaFold Server. The longest proteins that exceeded AlphaFold3 webserver limits were split into overlapping fragments of 3000 residues, which were then modeled individually. Proteins in which critical sections had low confidence (local pLDDT lower than 70) in AlphaFold3 prediction were additionally modeled using a local instance of ESMFold. We call all predictions collectively AF models.

All the models were subject to structure similarity searches using Dali^78^ and FATCAT ^49,50^ algorithms. To make structure similarity searches more accurate and easier to interpret, models of proteins longer than 600 residues were divided into fragments of 600 or fewer residues overlapping by 300 residues. The resulting models and models’ fragments were put as a searchable database on the public FATCAT^49^ web server. This database was then searched with queries representing different subcategories of the Outer membrane meander beta-barrels ECOD X class. This step identified additional beta-barrels which could not be found using BLASTP and HHPred (sequence-and HMM-based methods). We also retrieved the list of outer membrane beta-barrel proteins detected by the IsItABarrel program (https://isitabarrel.ku.edu/)^75^.

Independently, AF3 models of all *P. aeruginosa* proteins were analyzed by the in-house Python BarrelFinder program that searches for antiparallel beta sheets forming closed barrels and then fits an ellipsoid cylinder to the barrel^75^. BarrelFinder identified all meander beta barrels found in the previous steps, in addition to providing automated classification by the number of strands in the barrel as well as identification of loops connecting the barrel beta strands. While the periplasmic-facing loops are typically short (4-12 residues), some extracellular loops have lengths over 100 amino acids, forming “crowns”, which form primary immunogenic sites as well as participate in still mostly unidentified protein-protein interaction networks. Such crowns form a difficult to classify “psedo-domains”, regions with packing resembling that of single soluble protein domains but composed of several discontinuous segments (loops).

In the final stage of the analysis, all the collected 3D models of outer membrane beta barrels were divided into compact fragments corresponding to potential structural domains. This was initially done using an in-house program based on clustering of inter-residue contacts^79^ but then extensively corrected using PyMOL to visually inspect structures. These putative domains were then compared to structures of domains of experimentally characterized proteins retrieved from the ECOD database version 291^80^. (The ECOD version 291 released in April 2024 was the last ECOD release containing only experimental structures; we used its version clustered at 40% sequence identity available at ECOD website).

The structural comparisons were performed using the FATCAT program^49^ without allowing for twists in the aligned structures. The results of FATCAT searches were analyzed, and the putative domains were assigned to ECOD categories at the topology (T) level.

Finally, the sequences of the 105 *Pseudomonas* OMBBs were searched against proteins classified in the Transporter Classification Database (TCDB; https://tcdb.org/)^81^ using BLASTP^48^. For each OMBB, the TCDB family associated with the top-scoring BLASTP hit was assigned as its predicted TCDB family if the e-value was better than 0.05; otherwise, it was set to unknown.

### Extracellular-loop length analysis

The PA2590 cryo-EM structure was compared with AlphaFold3 models of 34 additional 22-stranded TonB-dependent transporters encoded by *P. aeruginosa* PAO1^40^. For each protein, the β-strands of the barrel were assigned from the secondary-structure topology, and the 11 extracellular loops were numbered sequentially from L1 to L11. Loop length was defined as the number of residues connecting the two β-strands flanking each extracellular loop, excluding residues assigned to the barrel strands. For each comparison protein (j), dissimilarity from PA2590 was quantified as the unweighted sum of the absolute differences between corresponding loop lengths:

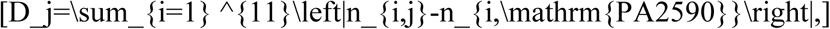

where (n_{i,j}) is the number of residues in loop (i) of protein (j). Scores were ranked in ascending order, with (D_j=0) indicating identical lengths across all 11 loops and lower values indicating greater similarity between loop-length profiles. A prominent surface-exposed loop pseudodomain (LPD) organization was evaluated independently by inspection of the three-dimensional models, based on spatially contiguous extracellular protrusions formed by adjacent loops, and was not included in the calculation of (D_j). Protein identities, UniProt accessions, individual loop lengths, LPD assignments, and calculated scores are provided in Supplementary file PA2590LoopAnalysis.xlsx.

### Recombinant protein cloning

The coding sequences (CDS) of the *P. aeruginosa* PAO1 outer membrane proteins were amplified by PCR from PAO1 genomic DNA using Phusion high-fidelity DNA polymerase (New England Biolabs, NEB) and the supplied GC buffer, to account for the high GC content of the PAO1 genome (∼60%). For all constructs, gene-specific primers (listed in Supplemental material, Table 3) were designed to amplify the CDS without the signal peptide. The native N-terminal signal peptide was omitted because the pTAMA-HIS-TEV expression vector carries an internal outer-membrane localization signal in the backbone. The pTAMA-HIS-TEV vector was linearized by restriction digestion with NcoI and BamHI. Due to poor recovery of the linearized vector after agarose gel excision, the digestion mixture, containing both the linearized pTAMA-HIS-TEV backbone and the original pfeA insert between the NcoI and BamHI sites, was used directly as the vector template in subsequent assembly reactions. Because the linearized vector retained sticky ends, one additional base was included in the 5′ overlap of the forward primers to maintain the correct reading frame at the NcoI junction. Insert fragments and the linearized pTAMA-HIS-TEV backbone were assembled using Gibson Assembly Master Mix (NEB, E2611) according to the manufacturer’s instructions. Assembly reactions (10 µL total) contained a 2:1 molar ratio of insert to vector DNA, 5 µL of 2× Gibson Assembly Master Mix, and nuclease-free water to volume, and were incubated at 50°C for 60 min in a thermal cycler. Following assembly, 1 µL of the reaction was transformed into NEB 5-alpha chemically competent *E. coli*. Cells were thawed on ice, mixed gently with the assembly product, incubated on ice for 30 min, heat-shocked at 42°C for 30 s, returned to ice for 2 min, and recovered in 950 µL LB medium at 37°C for 1 h with shaking at 175 rpm. Aliquots (200 µL) were plated on LB agar containing ampicillin and incubated overnight at 37°C. Individual colonies were screened by colony PCR using T7 sequencing primers and Taq DNA polymerase, and positive clones were confirmed by whole-plasmid sequencing. The rest of the clonal genes, including PA4579 and its mutants, were ordered from TWIST.

### Expression and purification

PA2590, PA3535, and wild-type (WT) PA4579 and its catalytic C256A variant were recombinantly expressed and purified from 3–6 L cultures using established laboratory protocols described previously^82,83^. PA2590 was purified by size-exclusion chromatography (SEC) in buffer containing 0.04% LDAO. Before SEC, PA3535 was reconstituted with amphipol A8-35 as described previously^82,83^. WT PA4579 underwent two sequential rounds of SEC, first in buffer containing *n*-dodecyl-β-D-maltoside (DDM) and then in buffer containing lauryl maltose neopentyl glycol (LMNG). Purified WT PA4579 and the C256A variant were obtained at final concentrations of 2.2 and 1.0 mg mL⁻¹, respectively, in the assay buffer described below.

### Cryo-EM grid preparation and data collection

The purified samples were applied to Quantifoil R 1.2/1.3 300 Mesh Gold grids. The grids were glow discharged for 90 s at 30 mA with a PELCO easiGlow™ Glow Discharge Cleaning System to obtain a hydrophilic surface. The glow-discharged grids were used to prepare vitrified samples with the Thermo Scientific Vitrobot Mark IV System. We applied 3 µl of purified protein to the glow-discharged surface of the grid at 4 °C, 100% humidity and blotted for 5.0s, with blot force 18.

The data were acquired with a 300 kV Titan Krios G3i microscope (Thermo Fisher) with a Falcon 4 detector at nominal magnification of 130,000× with a physical pixel size of 0.93985 Å. A phase plate was not used, and the objective aperture was not inserted. SerialEM was used for automated data collection in beam-image shift mode with 36 images (for PA2590) or 9 images (for PA3535) collected per stage movement, with a defocus range from -0.8 µm to -2.2 µm and beam-image shift compensation^84^. The slit width of the GIF Quantum Energy Filter was set to 25 eV for K3 and 10 eV for Falcon 4. Movies were dose-fractionated with a total dose of ∼50-70 e^-^/Å^2^.

### CryoEM data processing PA2590

7,424 movies were collected and imported into CryoSPARC followed by patch motion correction and patch CTF correction^85^. Particles were first picked using blob picker with diameter range of 70-130 Å on the first 1,000 micrographs. Picked particles were extracted with box size of 320 pixels Fourier cropped to 160 pixels, and cleaned by several rounds of 2D classifications. Classes showing clear protein features were used as templates for the next rounds of particle picking. 101 micrographs with the highest number of clean particles were selected for TOPAZ training^86^. 1,339,368 particles were then extracted using the trained model from all micrographs in 320-pixel boxes Fourier cropped to 160 pixels, followed by 2D classifications, resulting 479,854 cleaned particles.

Particles were sent to *ab initio* reconstruction job with five classes, resulting in one best class with 139,723 particles. After performing non-uniform refinement to confirm the quality of this class^87^, particles were re-extracted in 320-pixel boxes, resulting 138,753 final particles. These particles were further refined by non-uniform refinement to 3.13 Å. Due to the small size of the protein and the noise from the lipids surrounding the transmembrane region, local refinement using tight masks excluding the lipid molecules but not cutting through the protein density was conducted to improve resolution to 2.93 Å.

Particles from both datasets were then combined (257,624 total) and further cleaned by another round of *ab initio* reconstruction with two classes and heterogeneous refinement. A class of 151,313 particles showed high resolution features and was further refined by non-uniform refinement to a resolution of 3.92 Å^86^. Due to the small size of the protein and the noise from the lipids surrounding the transmembrane region, local refinement using tight masks excluding the lipid molecules but not cutting through the protein density was conducted to improve resolution to 3.57 Å.

### PA3535

6,516 movies were collected and processed with the standard motion correction and CTF estimation in CryoSPARC. 620 micrographs were processed initially to create templates for particle picking. Briefly, 894,508 particles were picked with a blob picker with a diameter range 50-130 Å. Particles were extracted in 256-pixel boxes and cleaned by 2D classification. Particles belonging to classes with clear protein features were selected and cleaned by another round of 2D classification, resulting in 147,551 particles. These particles were sent to *ab initio* reconstruction with one class and homogeneous refinement, producing 2.81 Å reconstruction.

2D templates were created from this 3D model, and 440,491 particles were picked by template-based particle picking from the original 620 micrographs. Particles were extracted in box size of 432 pixels, and 222,828 particles were retained after one round of 2D classification. 168 micrographs with the highest number of clean particles were selected for TOPAZ training^87^. 1,461,981 particles were then extracted using the trained model from all micrographs in 432-pixel boxes.

Several rounds of 2D classification and heterogeneous refinement using the initial 3D reconstruction and a decoy class created by junk particles were conducted to clean the extracted particles. Homogeneous refinement using only 448,807 retained clean particles resulted in a map with 2.51 Å resolution. However, the density of the transmembrane beta barrel is partially missing in this reconstruction due to motion between the soluble domain and the transmembrane domain. To achieve better reconstruction of each domain, separate local refinement was conducted as follows.

For the soluble domain reconstruction, local refinement was conducted with a mask surrounding only this domain without particle subtraction. The final resolution of the map was improved to 2.34 Å.

For the transmembrane domain reconstruction, due to its smaller size, heterogeneity introduced by the amphipol, and potential proteolytic cleavage at the linker between the soluble and transmembrane domain, simply masking around the barrel did result in resolution improvement in local reconstruction. To isolate particles with a rigid and intact transmembrane domain, 3D classification with soluble domain subtracted particles were performed. 178,786 particles with prominent densities at the transmembrane domain were selected from the 3D classification results and further refined by local refinement with masks surrounding the barrel, with the reconstruction achieving 3.23 Å resolution.

### Atomic model building

Initial atomic models were obtained by docking AlphaFold2 model of PA2590 and PA3535 to the maps using Molrep^37,88^. The model was then iteratively rebuilt manually in Coot and refined using *Phenix.real_space_refine* and Servalcat^89–95^. Model statistics can be found in Table 1.

**Table 1.**
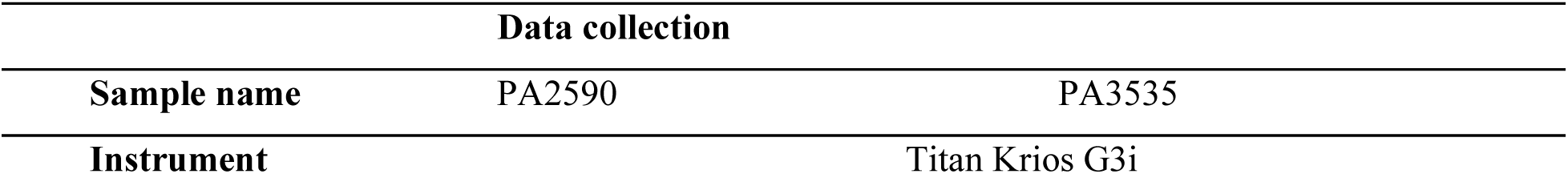
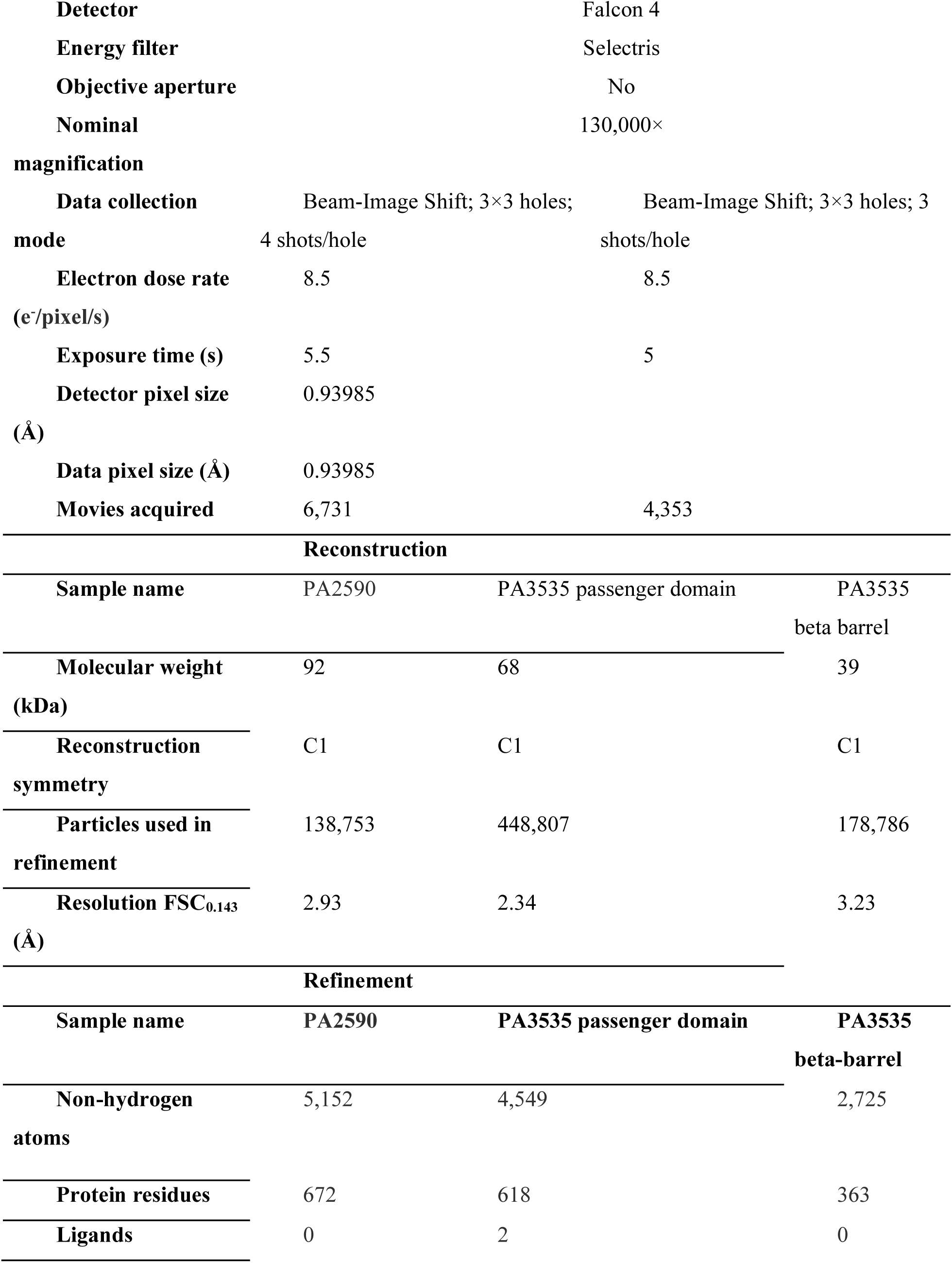
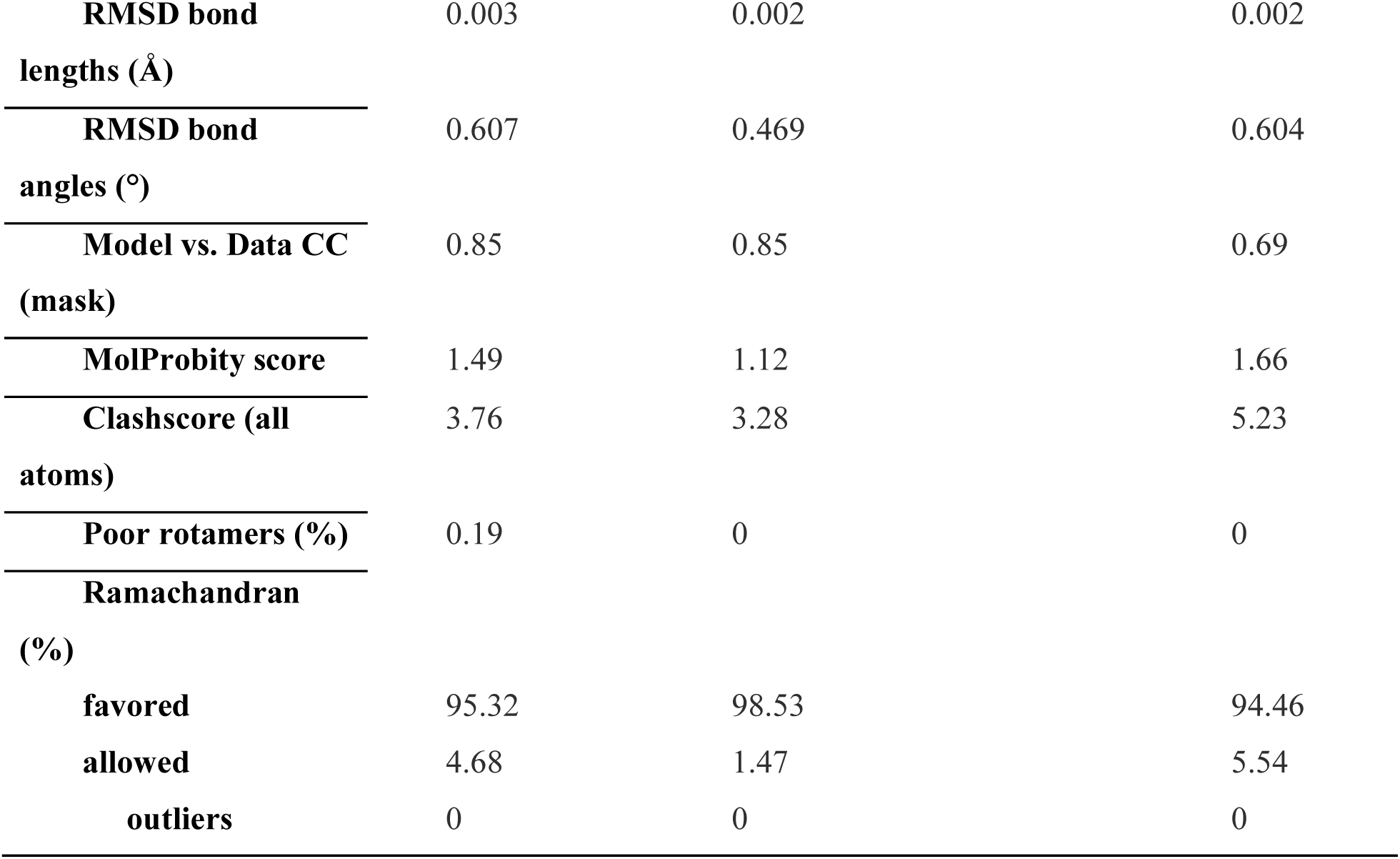
CryoEM data collection and processing.

### *In vitro* lipid cleavage reactions

PA4579 was expressed and purified using the laboratory standard protocol described above. For activity assays, PA4579 was TEV protease-cleaved and buffer-exchanged into assay buffer (Buffer F) and used at a concentration of 3 mg/ml. Buffer F consisted of 50 mM Tris-HCl, 150 mM NaCl, 0.5 mM TCEP, and 0.2% (w/v) LMNG, adjusted to pH 7.5. The lipopeptide substrate FSL-1 Biotin was prepared as a 5 mM stock in ultrapure water (ddH₂O). The fluorescent lipid substrate NBD-POPE (1-palmitoyl-2-{N-[7-nitro-2-1,3-benzoxadiazol-4-yl]amino}palmitoyl-sn-glycero-3-phosphoethanolamine) was stored as a 40 mg/mL primary stock in chloroform and diluted 10:1 in assay buffer to generate a 4 mg/mL working stock. Four reaction conditions were prepared in a total volume of 400 µL each: (i) WT PA4579 with FSL-1 Biotin and NBD-POPE, (ii) C256A mutant with FSL-1 Biotin and NBD-POPE, (iii) FSL-1 Biotin with NBD-POPE (no enzyme control), and (iv) NBD-POPE alone (substrate-only control). Enzyme concentrations were standardized to 800 nM (approximately 0.05 mg/mL) across all enzyme-containing reactions. Each reaction containing NBD-POPE received 4 µL of the 4 mg/mL working stock (16 µg total phosphatidylethanolamine per reaction), and FSL-1 Biotin-containing reactions received 2 µL of the 5 mM stock. Reaction volumes were brought to 400 µL with assay buffer. Following addition of NBD-POPE but prior to enzyme addition, all reaction mixtures were sonicated in a water bath sonicator until optically clear (approximately 15 min, with intermittent vortexing every ∼5 min) to ensure complete lipid suspension. Enzyme was then added to the appropriate reactions, and all samples were incubated at 37 °C for approximately 48 h.

### Lipid extraction and high-performance thin-layer chromatography (HPTLC)

Lipid PE standards 1-palmitoyl-2-oleoyl-sn-glycero-3-phosphoethanolamine (16:0-18:1 PE (POPE); #A85757), 2-dioleoyl-sn-glycero-3-phosphoethanolamine (18:1 PE (DOPE); #A80725), and 1-oleoyl-2-{12-[(7-nitro-2-1,3-benzoxadiazol-4-yl)amino] dodecanoyl}-sn-glycero-3-phosphoethanolamine (18:1-12:0 NBD-PE (NBD-OLPE); #A81156) were obtained from Avanti Polar Lipids. Silica gel 60 HPTLC plates (Supelco, #105641) and all HPLC-grade solvents were purchased from EMD Millipore. Lipids were extracted using the Bligh and Dyer methanol/chloroform extraction method^96^ with modifications^97^. The chloroform phase was collected and dried under nitrogen, and lipid films were reconstituted in 100 μL chloroform: methanol (2:1, v/v). Silica gel 60 HPTLC plates were pre-washed with methanol: ethyl acetate (6:4, v/v) and activated at 110 °C for 30 min. Lipid extracts were applied onto the HPTLC plate using a LINOMAT 5 automated sample applicator (CAMAG, Switzerland). Phospholipids were resolved by automated multiple development using a CAMAG AMD2 system (CAMAG, Switzerland), and PE, LPE, and NBD-labelled phosphatidylethanolamine (NBD-PE) standards were included on each plate for analysis. Plates were developed over 90 mm with dichloromethane acetate (80:16:4, v/v/v), followed by a second 90 mm development using chloroform acetate: acetone: isopropanol: ethanol: methanol: water: acid (30:6:6:6:16:28:6:2, v/v/v/v/v/v/v/v)^97^.

Following chromatographic separation, NBD-labelled lipids were first detected by fluorescence imaging (excitation/emission, 465/536 nm), allowing selective visualization of NBD-PE and its hydrolytic product, NBD-lysophosphatidylethanolamine (NBD-LPE), thereby enabling direct detection of PE-to-LPE conversion. The same HPTLC plates were subsequently immersed in 10% (w/v) CuSO₄ in 8% (v/v) H₃PO₄, charred for 8-10 min at 168°C and imaged using an iBright FL1000 imaging system (Thermo Fisher Scientific) with excitation/emission wavelengths of 540/590 nm to visualize total phospholipid species (Supplemental Figures 4-6).

## Supporting information

Suppl_Table_SX; Suppl_PA2590LoopAnalysis ; 4579Gels added_Supplement figs and tables

## Figure preparation

For visualization of the experimental structures in PyMOL^98^ and ChimeraX^99^, the DSSP command was used to assign secondary-structure elements consistently, enabling accurate representation of β-strands, α-helices, and loop regions.

## Data availability

The PA2590 atomic coordinate (PDB 38PE) and map (EMD-79031), and the PA3535 atomic coordinates (beta barrel: PDB 38PG; passenger domain: PDB 38PF) and maps (beta barrel: EMD-79034; passenger domain: EMD-79033; composite map: EMD-79035; consensus map: EMD-79032) have been deposited in the Protein Data Bank (http://wwpdb.org/) and the Electron Microscopy Data Bank (https://www.ebi.ac.uk/emdb/), respectively.

## Acknowledgments

This work was supported by the Cumming Medical Research Fund (CMRF) at the University of Calgary and, in part, by federal funds from the National Institute of Allergy and Infectious Diseases, National Institutes of Health, Department of Health and Human Services, under Contract No. 75N93022C00035 (Center for Structural Biology of Infectious Diseases, CSBID) to A.S. and E.K.

Additional funding and technical expertise for bacterial antigen production were provided through the University of Calgary Immunotherapeutics Core Facility.

We thank the staff of the Cryo-Electron Microscopy Facility (CEMF) at UT Southwestern Medical Center for maintaining the electron microscopes used in this study. The CEMF receives support from the Cancer Prevention & Research Institute of Texas (CPRIT) through grants RP170644 and RP220582.

TR acknowledges funding from Mount Royal University FST Innovation Grant No.104891.

Computations were performed using the computer clusters and data storage resources of the UCR High Performance Computer Cluster (HPCC), which were funded by NSF (MRI-2215705, MRI-1429826) and NIH (1S10OD016290-01A1) grants.

## Author contributions

E.K.: Conceptualization, project administration, supervision, investigation, methodology, formal analysis, visualization, writing – original draft, and writing – review and editing. L.J.: Conceptualization, formal analysis, methodology, visualization, writing – original draft, and writing – review and editing. J.T.: Investigation and writing – original draft, writing – review and editing. D.P.: Investigation, visualization, and writing – original draft. Y.G.: Investigation, formal analysis, and visualization; writing – review and editing. T.R.: Investigation, funding acquisition, formal analysis, visualization, writing – original draft, and writing – review and editing. D.B.: Formal analysis, funding acquisition, and writing – review and editing. J.J.H.: Formal analysis and writing – review and editing. A.G.: Formal analysis, methodology, supervision, and writing – review and editing. A.S.: Conceptualization, formal analysis; funding acquisition, supervision, and writing – review and editing.

## Code availability, if applicable

## References

1. Reynolds, D. & Kollef, M. The Epidemiology and Pathogenesis and Treatment of Pseudomonas aeruginosa Infections: An Update. Drugs 81, 2117–2131 (2021).

2. Sati, H. et al. The WHO Bacterial Priority Pathogens List 2024: a prioritisation study to guide research, development, and public health strategies against antimicrobial resistance. Lancet Infect Dis 25, 1033–1043 (2025).

3. Diggle, S.P. & Whiteley, M. Microbe Profile: Pseudomonas aeruginosa: opportunistic pathogen and lab rat. Microbiology (Reading) 166, 30–33 (2020).

4. Gellatly, S.L. & Hancock, R.E. Pseudomonas aeruginosa: new insights into pathogenesis and host defenses. Pathog Dis 67, 159–73 (2013).

5. Qin, S. et al. Pseudomonas aeruginosa: pathogenesis, virulence factors, antibiotic resistance, interaction with host, technology advances and emerging therapeutics. Signal Transduct Target Ther 7, 199 (2022).

6. Mulcahy, L.R., Isabella, V.M. & Lewis, K. Pseudomonas aeruginosa biofilms in disease. Microb Ecol 68, 1–12 (2014).

7. Maurice, N.M., Bedi, B. & Sadikot, R.T. Pseudomonas aeruginosa Biofilms: Host Response and Clinical Implications in Lung Infections. Am J Respir Cell Mol Biol 58, 428–439 (2018).

8. Ude, J. et al. Outer membrane permeability: Antimicrobials and diverse nutrients bypass porins in Pseudomonas aeruginosa. Proc Natl Acad Sci U S A 118(2021).

9. Hancock, R.E. & Brinkman, F.S. Function of pseudomonas porins in uptake and efflux. Annu Rev Microbiol 56, 17–38 (2002).

10. Pagès, J.M., James, C.E. & Winterhalter, M. The porin and the permeating antibiotic: a selective diffusion barrier in Gram-negative bacteria. Nat Rev Microbiol 6, 893–903 (2008).

11. Henderson, J.C. et al. The Power of Asymmetry: Architecture and Assembly of the Gram-Negative Outer Membrane Lipid Bilayer. Annu Rev Microbiol 70, 255–78 (2016).

12. Sandoz, K.M. et al. β-Barrel proteins tether the outer membrane in many Gram-negative bacteria. Nat Microbiol 6, 19–26 (2021).

13. Chevalier, S. et al. Structure, function and regulation of Pseudomonas aeruginosa porins. FEMS Microbiol Rev 41, 698–722 (2017).

14. Fito-Boncompte, L. et al. Full virulence of Pseudomonas aeruginosa requires OprF. Infect Immun 79, 1176–86 (2011).

15. Li, H., Luo, Y.F., Williams, B.J., Blackwell, T.S. & Xie, C.M. Structure and function of OprD protein in Pseudomonas aeruginosa: from antibiotic resistance to novel therapies. Int J Med Microbiol 302, 63–8 (2012).

16. Stover, C.K. et al. Complete genome sequence of Pseudomonas aeruginosa PAO1, an opportunistic pathogen. Nature 406, 959–64 (2000).

17. Cornelis, P. & Bodilis, J. A survey of TonB-dependent receptors in fluorescent pseudomonads. Environ Microbiol Rep 1, 256–62 (2009).

18. Winsor, G.L. et al. Enhanced annotations and features for comparing thousands of Pseudomonas genomes in the Pseudomonas genome database. Nucleic Acids Res 44, D646–53 (2016).

19. Freschi, L. et al. The Pseudomonas aeruginosa Pan-Genome Provides New Insights on Its Population Structure, Horizontal Gene Transfer, and Pathogenicity. Genome Biol Evol 11, 109–120 (2019).

20. Lee, A.J. et al. Compendium-Wide Analysis of Pseudomonas aeruginosa Core and Accessory Genes Reveals Transcriptional Patterns across Strains PAO1 and PA14. mSystems 8, e0034222 (2023).

21. Botelho, J. et al. Phylogroup-specific variation shapes the clustering of antimicrobial resistance genes and defence systems across regions of genome plasticity in Pseudomonas aeruginosa. EBioMedicine 90, 104532 (2023).

22. Blonder, J. et al. Global analysis of the membrane subproteome of Pseudomonas aeruginosa using liquid chromatography-tandem mass spectrometry. J Proteome Res 3, 434–44 (2004).

23. Duchesne, R. et al. A proteomic approach of SigX function in Pseudomonas aeruginosa outer membrane composition. J Proteomics 94, 451–9 (2013).

24. Tamber, S., Ochs, M.M. & Hancock, R.E. Role of the novel OprD family of porins in nutrient uptake in Pseudomonas aeruginosa. J Bacteriol 188, 45–54 (2006).

25. Eren, E. et al. Substrate specificity within a family of outer membrane carboxylate channels. PLoS Biol 10, e1001242 (2012).

26. Bhamidimarri, S.P. et al. Acquisition of ionic copper by the bacterial outer membrane protein OprC through a novel binding site. PLoS Biol 19, e3001446 (2021).

27. Chan, D.C.K. & Burrows, L.L. Pseudomonas aeruginosa FpvB Is a High-Affinity Transporter for Xenosiderophores Ferrichrome and Ferrioxamine B. mBio 14, e0314922 (2023).

28. Knejski, P.P., Erramilli, S.K. & Kossiakoff, A.A. Chaperone-assisted cryo-EM structure of P. aeruginosa PhuR reveals molecular basis for heme binding. Structure 32, 411–423.e6 (2024).

29. Benedens, M. et al. Assembly and the gating mechanism of the Pel exopolysaccharide export complex PelBC of Pseudomonas aeruginosa. Nat Commun 16, 5249 (2025).

30. Tralau, T. et al. Transcriptomic analysis of the sulfate starvation response of Pseudomonas aeruginosa. J Bacteriol 189, 6743–50 (2007).

31. Noinaj, N., Guillier, M., Barnard, T.J. & Buchanan, S.K. TonB-dependent transporters: regulation, structure, and function. Annu Rev Microbiol 64, 43–60 (2010).

32. Fujita, M. et al. A TonB-dependent receptor constitutes the outer membrane transport system for a lignin-derived aromatic compound. Commun Biol 2, 432 (2019).

33. Silale, A. & van den Berg, B. TonB-Dependent Transport Across the Bacterial Outer Membrane. Annu Rev Microbiol 77, 67–88 (2023).

34. McPhee, J.B. et al. The major outer membrane protein OprG of Pseudomonas aeruginosa contributes to cytotoxicity and forms an anaerobically regulated, cation-selective channel. FEMS Microbiol Lett 296, 241–7 (2009).

35. Kucharska, I., Liang, B., Ursini, N. & Tamm, L.K. Molecular Interactions of Lipopolysaccharide with an Outer Membrane Protein from Pseudomonas aeruginosa Probed by Solution NMR. Biochemistry 55, 5061–72 (2016).

36. Edrington, T.C., Kintz, E., Goldberg, J.B. & Tamm, L.K. Structural basis for the interaction of lipopolysaccharide with outer membrane protein H (OprH) from Pseudomonas aeruginosa. J Biol Chem 286, 39211–23 (2011).

37. Jumper, J. et al. Highly accurate protein structure prediction with AlphaFold. Nature 596, 583–589 (2021).

38. Baek, M. et al. Accurate prediction of protein structures and interactions using a three-track neural network. Science 373, 871–876 (2021).

39. UniProt: the Universal Protein Knowledgebase in 2025. Nucleic Acids Res 53, D609–d617 (2025).

40. Abramson, J. et al. Accurate structure prediction of biomolecular interactions with AlphaFold 3. Nature 630, 493–500 (2024).

41. Lin, Z. et al. Evolutionary-scale prediction of atomic-level protein structure with a language model. Science 379, 1123–1130 (2023).

42. Altschul, S.F., Gish, W., Miller, W., Myers, E.W. & Lipman, D.J. Basic local alignment search tool. J Mol Biol 215, 403–10 (1990).

43. Zimmermann, L. et al. A Completely Reimplemented MPI Bioinformatics Toolkit with a New HHpred Server at its Core. J Mol Biol 430, 2237–2243 (2018).

44. Ye, Y. & Godzik, A. FATCAT: a web server for flexible structure comparison and structure similarity searching. Nucleic Acids Res 32, W582–5 (2004).

45. Ernst, R.K. et al. The Pseudomonas aeruginosa lipid A deacylase: selection for expression and loss within the cystic fibrosis airway. J Bacteriol 188, 191–201 (2006).

46. Rutten, L. et al. Crystal structure and catalytic mechanism of the LPS 3-O-deacylase PagL from Pseudomonas aeruginosa. Proc Natl Acad Sci U S A 103, 7071–6 (2006).

47. Luckett, J.C. et al. A novel virulence strategy for Pseudomonas aeruginosa mediated by an autotransporter with arginine-specific aminopeptidase activity. PLoS Pathog 8, e1002854 (2012).

48. Altschul, S.F. et al. Gapped BLAST and PSI-BLAST: a new generation of protein database search programs. Nucleic Acids Res 25, 3389–402 (1997).

49. Ye, Y. & Godzik, A. Flexible structure alignment by chaining aligned fragment pairs allowing twists. Bioinformatics 19 **Suppl 2**, ii246–55 (2003).

50. Li, Z., Jaroszewski, L., Iyer, M., Sedova, M. & Godzik, A. FATCAT 2.0: towards a better understanding of the structural diversity of proteins. Nucleic Acids Res 48, W60–w64 (2020).

51. Premsuriya, J., Mosbahi, K., Atanaskovic, I., Kleanthous, C. & Walker, D. Outer membrane translocation of pyocins via the copper regulated TonB-dependent transporter CrtA. Biochem J 480, 1035–1049 (2023).

52. Zhou, H. et al. Spatial transcriptomics identifies novel Pseudomonas aeruginosa virulence factors. Cell Genom 5, 100805 (2025).

53. Henderson, I.R., Navarro-Garcia, F., Desvaux, M., Fernandez, R.C. & Ala’Aldeen, D. Type V protein secretion pathway: the autotransporter story. Microbiol Mol Biol Rev 68, 692–744 (2004).

54. Clarke, K.R. et al. Phylogenetic Classification and Functional Review of Autotransporters. Front Immunol 13, 921272 (2022).

55. Kida, Y., Taira, J., Yamamoto, T., Higashimoto, Y. & Kuwano, K. EprS, an autotransporter protein of Pseudomonas aeruginosa, possessing serine protease activity induces inflammatory responses through protease-activated receptors. Cell Microbiol 15, 1168–81 (2013).

56. Wilhelm, S., Gdynia, A., Tielen, P., Rosenau, F. & Jaeger, K.E. The autotransporter esterase EstA of Pseudomonas aeruginosa is required for rhamnolipid production, cell motility, and biofilm formation. J Bacteriol 189, 6695–703 (2007).

57. van den Berg, B. Crystal structure of a full-length autotransporter. J Mol Biol 396, 627–33 (2010).

58. Arachchige, E.J., Rahman, M.S., Singendonk, K.S. & Kim, K.H. Structural and Functional Characterization of Pseudomonas aeruginosa Virulence Factor AaaA, an Autotransporter with Arginine-Specific Aminopeptidase Activity. J Mol Biol 437, 169358 (2025).

59. Reddy, B.L. & Saier, M.H., Jr. Properties and Phylogeny of 76 Families of Bacterial and Eukaryotic Organellar Outer Membrane Pore-Forming Proteins. PLoS One 11, e0152733 (2016).

60. Blum, M. et al. InterPro: the protein sequence classification resource in 2025. Nucleic Acids Res 53, D444–d456 (2025).

61. Schaeffer, R.D. et al. ECOD: integrating classifications of protein domains from experimental and predicted structures. Nucleic Acids Res 53, D411–d418 (2025).

62. Blum, M. et al. The InterPro protein families and domains database: 20 years on. Nucleic Acids Res 49, D344–d354 (2021).

63. Kida, Y., Taira, J. & Kuwano, K. EprS, an autotransporter serine protease, plays an important role in various pathogenic phenotypes of Pseudomonas aeruginosa. Microbiology (Reading) 162, 318–329 (2016).

64. Hor, L. et al. Crystal structure of a subtilisin-like autotransporter passenger domain reveals insights into its cytotoxic function. Nat Commun 14, 1163 (2023).

65. Leyton, D.L. et al. Size and conformation limits to secretion of disulfide-bonded loops in autotransporter proteins. J Biol Chem 286, 42283–42291 (2011).

66. Oomen, C.J. et al. Structure of the translocator domain of a bacterial autotransporter. Embo j 23, 1257–66 (2004).

67. Ekici, O.D., Paetzel, M. & Dalbey, R.E. Unconventional serine proteases: variations on the catalytic Ser/His/Asp triad configuration. Protein Sci 17, 2023–37 (2008).

68. van Kempen, M. et al. Fast and accurate protein structure search with Foldseek. Nat Biotechnol (2023).

69. Bryan, P., Pantoliano, M.W., Quill, S.G., Hsiao, H.Y. & Poulos, T. Site-directed mutagenesis and the role of the oxyanion hole in subtilisin. Proc Natl Acad Sci U S A 83, 3743–5 (1986).

70. Armbruster, K.M. et al. Identification and characterization of the lipoprotein N-acyltransferase in Bacteroides. Proc Natl Acad Sci U S A 121, e2410909121 (2024).

71. Akama, H. et al. Crystal structure of the drug discharge outer membrane protein, OprM, of Pseudomonas aeruginosa: dual modes of membrane anchoring and occluded cavity end. J Biol Chem 279, 52816–9 (2004).

72. Hoang, H.H. et al. Outer membrane targeting of Pseudomonas aeruginosa proteins shows variable dependence on the components of Bam and Lol machineries. mBio 2(2011).

73. Takeda, K. et al. Crystal structures of bacterial lipoprotein localization factors, LolA and LolB. Embo j 22, 3199–209 (2003).

74. Taniguchi, N., Matsuyama, S. & Tokuda, H. Mechanisms underlying energy-independent transfer of lipoproteins from LolA to LolB, which have similar unclosed {beta}-barrel structures. J Biol Chem 280, 34481–8 (2005).

75. Montezano, D., Bernstein, R., Copeland, M.M. & Slusky, J.S.G. General features of transmembrane beta barrels from a large database. Proc Natl Acad Sci U S A 120, e2220762120 (2023).

76. Chandrasekaran, P., Weiskirchen, S. & Weiskirchen, R. Structure, Functions, and Implications of Selected Lipocalins in Human Disease. Int J Mol Sci 25(2024).

77. Moynié, L. et al. The complex of ferric-enterobactin with its transporter from Pseudomonas aeruginosa suggests a two-site model. Nat Commun 10, 3673 (2019).

78. Holm, L. & Sander, C. Protein structure comparison by alignment of distance matrices. J Mol Biol 233, 123–38 (1993).

79. Feldman, H.J. Identifying structural domains of proteins using clustering. BMC Bioinformatics 13, 286 (2012).

80. Cheng, H. et al. ECOD: an evolutionary classification of protein domains. PLoS Comput Biol 10, e1003926 (2014).

81. Saier, M.H. et al. The Transporter Classification Database (TCDB): 2021 update. Nucleic Acids Res 49, D461–d467 (2021).

82. Patel, D.H. et al. Cryo-EM SPR structures of Salmonella typhimurium ArnC; the key enzyme in lipid-A modification conferring polymyxin resistance. Protein Sci 34, e70037 (2025).

83. Karimullina, E. et al. Structural architecture of TolQ-TolR inner membrane protein complex from opportunistic pathogen Acinetobacter baumannii. Sci Adv 11, eadq9845 (2025).

84. Mastronarde, D.N. Automated electron microscope tomography using robust prediction of specimen movements. J Struct Biol 152, 36–51 (2005).

85. Punjani, A., Rubinstein, J.L., Fleet, D.J. & Brubaker, M.A. cryoSPARC: algorithms for rapid unsupervised cryo-EM structure determination. Nat Methods 14, 290–296 (2017).

86. Bepler, T. et al. Positive-unlabeled convolutional neural networks for particle picking in cryo-electron micrographs. Nat Methods 16, 1153–1160 (2019).

87. Punjani, A., Zhang, H. & Fleet, D.J. Non-uniform refinement: adaptive regularization improves single-particle cryo-EM reconstruction. Nat Methods 17, 1214–1221 (2020).

88. Vagin, A. & Teplyakov, A. Molecular replacement with MOLREP. Acta Crystallogr D Biol Crystallogr 66, 22–5 (2010).

89. Emsley, P. & Cowtan, K. Coot: model-building tools for molecular graphics. Acta Crystallogr D Biol Crystallogr 60, 2126–32 (2004).

90. Krissinel, E. & Henrick, K. Secondary-structure matching (SSM), a new tool for fast protein structure alignment in three dimensions. Acta Crystallogr D Biol Crystallogr 60, 2256–68 (2004).

91. Liebschner, D. et al. Macromolecular structure determination using X-rays, neutrons and electrons: recent developments in Phenix. Acta Crystallogr D Struct Biol 75, 861–877 (2019).

92. Murshudov, G.N. et al. REFMAC5 for the refinement of macromolecular crystal structures. Acta Crystallogr D Biol Crystallogr 67, 355–67 (2011).

93. Yamashita, K., Palmer, C.M., Burnley, T. & Murshudov, G.N. Cryo-EM single-particle structure refinement and map calculation using Servalcat. Acta Crystallogr D Struct Biol 77, 1282–1291 (2021).

94. Emsley, P., Lohkamp, B., Scott, W.G. & Cowtan, K. Features and development of Coot. Acta Crystallogr D Biol Crystallogr 66, 486–501 (2010).

95. Afonine, P.V. et al. Real-space refinement in PHENIX for cryo-EM and crystallography. Acta Crystallogr D Struct Biol 74, 531–544 (2018).

96. Bligh, E.G. & Dyer, W.J. A rapid method of total lipid extraction and purification. Can J Biochem Physiol 37, 911–7 (1959).

97. Churchward, M.A., Brandman, D.M., Rogasevskaia, T. & Coorssen, J.R. Copper (II) sulfate charring for high sensitivity on-plate fluorescent detection of lipids and sterols: quantitative analyses of the composition of functional secretory vesicles. J Chem Biol 1, 79–87 (2008).

98. Schrödinger, L.L.C. The PyMOL Molecular Graphics System, Version 2.3. (2020).

99. Meng, E.C. et al. UCSF ChimeraX: Tools for structure building and analysis. Protein Sci 32, e4792 (2023).

