## Supplementary material for "Charting the Expansive Outer Membrane β-Barrel Landscape in *Pseudomonas aeruginosa*": Suppl_Table_SX; Suppl_PA2590LoopAnalysis ; 4579Gels added_Supplement figs and tables: 4579Gels added_Supplement figs and tables file copy.docx

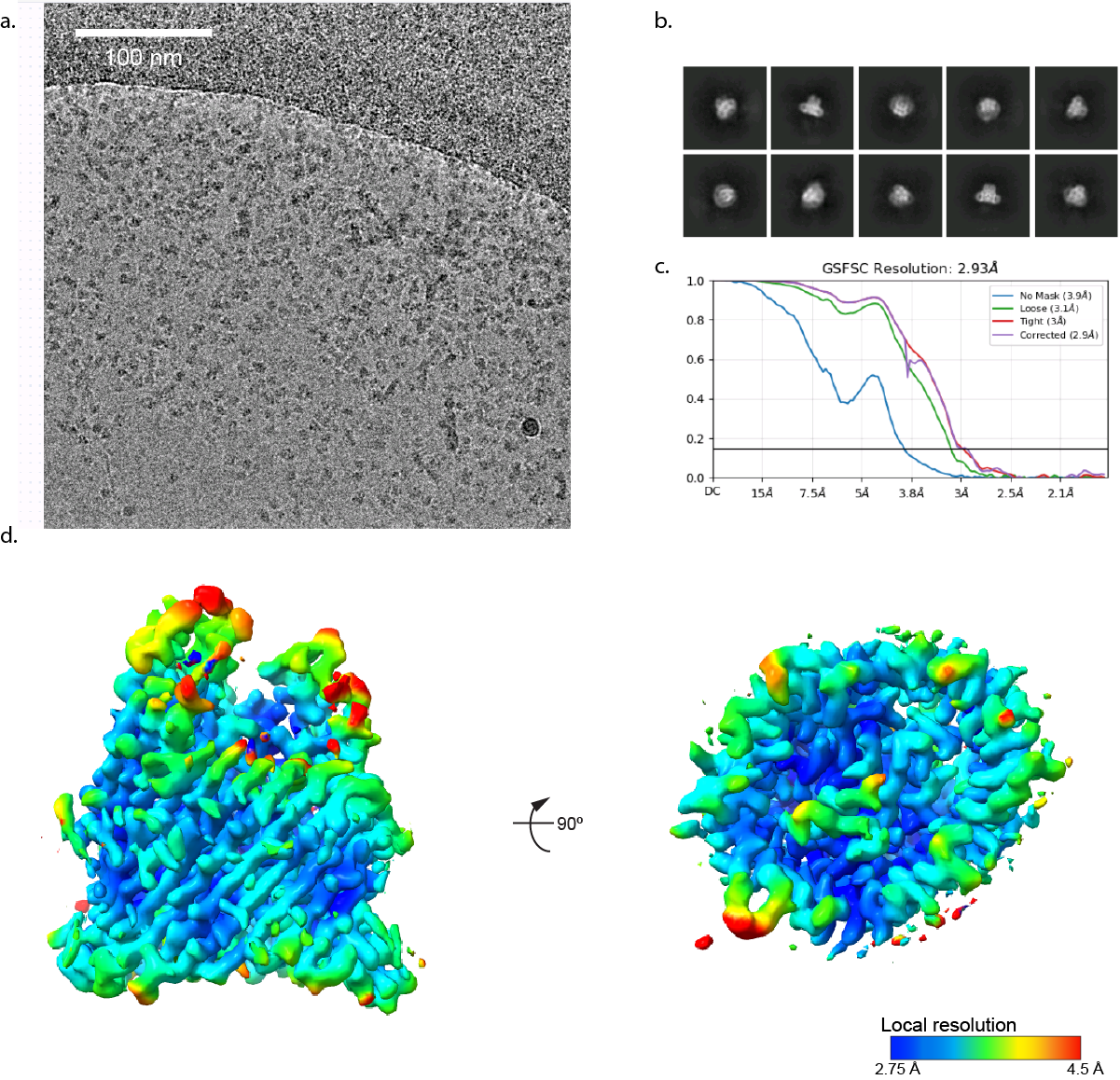

Supplementary Figure 1 Chart of PA2590 single-particle reconstruction.

Supplementary table 1: Functional annotation of predicted 22-stranded TonB-dependent outer-membrane β-barrel proteins in Pseudomonas aeruginosa PAO1. The complete predicted domain architectures of these proteins are available in the Pathogens3D Pseudomonas porins dataset: <https://proteins.pathogens3d.org/dataset/pseudomonas-porins>.

| **PA number** | **UniProt accession** | **Best current name** | **Functional assignment from literature** | **PMID(s)** | **Homology note / non-Pseudomonas analog** |
| --- | --- | --- | --- | --- | --- |
| PA4897 | Q9HUR6 | OptI / PA4897 CSS transducer | Cell-surface-signaling-associated TonB-dependent transducer; cognate ligand/substrate not identified. Older annotation suggested heme-receptor-like homology, but no receptor-specific transport paper was found. | PMID: 18086184; 29555629 | KEGG assigns PA4897 to K16087, hemoglobin/transferrin/lactoferrin receptor protein, but no experimentally confirmed ligand or transport substrate is available. |
| PA2070 | Q9I245 | OptM | Poorly characterized receptor/transducer linked to ECF/CSS regulatory screens; no substrate identified. | PMID: 18086184; 29555629 |  |
| PA2057 | Q9I258 | SppR | Part of the SppR-SppABCD system; functional work links the locus to swarming modulation. Proposed xenosiderophore-type role remains unconfirmed. | PMID: 26995781 |  |
| PA0192 | Q9I6U2 | OptP | Predicted TBDR historically annotated by homology to FyuA/pesticin-receptor-like proteins; no ligand/function assigned. | PMID: 29555629 | FyuA/pesticin-receptor-like homology noted; 28% amino-acid identity to pathogenic *E. coli* FyuA |
| PA4514 | G3XCY8 | PiuA | Siderophore-drug entry receptor. PiuA/PirA-family receptors are implicated in uptake of BAL30072, MC-1 and cefiderocol-like siderophore conjugates. Natural physiological siderophore ligand remains unresolved. | PMID: 28137795; 29555629 | *P. aeruginosa* functional/structural evidence; 60% amino-acid identity to *E. coli* Fiu P75780. |
| PA4168 | Q9HWL3 | FpvB | Alternative type I ferripyoverdine receptor; newer work supports high-affinity uptake of ferrichrome and ferrioxamine B with relatively poor pyoverdine transport. | PMID: 15184553; 36507834 | *P. aeruginosa* functional evidence |
| PA2466 | Q9I116 | FoxA | Ferrioxamine B/nocardamine receptor and cell-surface-signaling transducer; structural work supports ferrioxamine B transport and TonB engagement. | PMID: 16484199; 31385808; 32902248 | *P. aeruginosa* functional/structural evidence; 43-47% amino-acid identity to *Salmonella enterica* serovar Typhimurium, *Yersinia enterocolitica*, and *Erwinia amylovora* |
| PA1910 | Q9I2J4 | FemA | Receptor/transducer in the FemI-FemR-FemA CSS system; responds to mycobactin/carboxymycobactin-type mycobacterial siderophores and supports their utilization; aeruginoic and dihydroaeruginoic acids iron complexes are transported via FemA | PMID: 18086184;  40035455 | *P. aeruginosa* functional evidence; mycobactin/carboxymycobactin are non-*Pseudomonas* ligands, but receptor is characterized in *P. aeruginosa;* |
| PA0470 | Q9I648 | FiuA | Ferrichrome/desferrichrome receptor; also implicated in biofilm and virulence-associated phenotypes. | PMID: 16484199; 20047910; 27190289 | *P. aeruginosa* functional evidence; 32% amino-acid identity to *E. coli* FhuA P06971 |
| PA0434 | Q9I683 | CrtA / OptJ | Copper-responsive TBDR and receptor exploited by S-type pyocins SX1/SX2; also identified as a siderophore-drug susceptibility hit. Natural transported substrate remains unclear. | PMID: 29555629;  37399084;  41570985 | *P. aeruginosa* pyocin/copper-responsive evidence |
| PA4221 | P42512 | FptA | Canonical ferric-pyochelin outer-membrane receptor; also emerges in siderophore-drug conjugate uptake screens. | PMID: 8288523; 16139844; 29555629 | *P. aeruginosa* functional and structural evidence |
| PA2398 | P48632 | FpvA | Major ferripyoverdine receptor. Mediates pyoverdine-Fe uptake and participates in pyoverdine-dependent cell-surface signaling through the FpvR/FpvI/FpvA axis. | PMID: 8335619; 29555629 | *P. aeruginosa* functional evidence |
| PA4837 | Q9HUX3 | CntO | Pseudopaline metallophore receptor involved in Zn and Ni acquisition under metal limitation. | PMID: 26290475; 29214991; 41532760 | *P. aeruginosa* functional evidence |
| PA3901 | Q9HXB2 | FecA | Ferric-citrate receptor required for citrate-promoted growth and citrate-mediated iron uptake. | PMID: 19118371 | *P. aeruginosa* functional evidence; 64% amino-acid identity to *E. coli* FecA P13036 |
| PA2911 | Q9HZT6 | OptE | Zinc-restriction-associated TonB-dependent outer-membrane protein; also a siderophore-drug conjugate susceptibility hit. Natural ligand is not well defined. | PMID: 26290475; 29555629 | *P. aeruginosa* zinc-homeostasis evidence; no defined substrate analog |
| PA2590 | Q9I0P7 | OptF | Poorly characterized TBDR. Appears in receptor-repertoire/CSS-associated screens, but no dedicated transport or substrate paper was found. | PMID: 18086184; 29555629 | No confirmed *P. aeruginosa* ligand |
| PA2089 | Q9I226 | OptL | Poorly characterized CSS-associated/predicted receptor; no substrate identified. | PMID: 18086184; 29555629 | No confirmed ligand |
| PA1322 | Q9I422 | PfuA | Predicted ferrichrome-like receptor; identified as a siderophore-drug conjugate susceptibility hit. Natural substrate remains unresolved. | PMID: 29555629 | Predicted ferrichrome-like/siderophore-drug receptor. |
| PA0151 | Q9I6Y1 | PupB-like | CSS-associated predicted receptor, PupB-like by homology. No experimentally assigned ligand found. | PMID: 18086184; 29555629 | PupB-like homology noted |
| PA1613 | Q9I3A9 | CirA-like | Predicted CirA-like receptor; no assigned substrate/function found. | PMID: 29555629 | 35% amino-acid identity to *E. coli* CirA P17315. |
| PA2335 | Q9I1E3 | OptO | Predicted TBDR; no receptor-specific substrate/function paper found. | PMID: 29555629 |  |
| PA2688 | Q05098 | PfeA | Catecholate siderophore receptor, especially ferrienterobactin and synthetic catecholate siderophores; overlaps with PirA. | PMID: 15899402; 31413254 | *P. aeruginosa* functional/structural evidence; 60% amino-acid identity to *E. coli* FepA P05825. |
| PA0931 | Q9I527 | PirA | Second ferrienterobactin/catecholate receptor; also involved in siderophore-drug entry with PiuA-like systems. | PMID: 15899402; 28137795 | *P. aeruginosa* functional/structural evidence; 56% amino-acid identity to *E. coli* FepA P05825. |
| PA4675 | Q9HVC0 | ChtA | Xenosiderophore receptor most strongly linked to aerobactin-Fe; also implicated in schizokinen/rhizobactin-class citrate-hydroxamate siderophores, with later work refining substrate specificity. | PMID: 16549659;  37758521 | *P. aeruginosa* functional evidence; 46% amino-acid identity to *E. coli* IutA. |
| PA4156 | Q9HWM5 | FvbA | Vibriobactin receptor enabling use of the Vibrio-derived catecholate siderophore vibriobactin. | PMID: 21546589 | *P. aeruginosa* functional evidence for non-Pseudomonas ligand |
| PA1302 | Q9I442 | HxuA | Heme-associated CSS receptor/transducer in the Hxu system. Major role appears to be signaling; minor heme-acquisition role when HasR/PhuR or other iron sources are available. | PMID: 31390127; 36169312 | *P. aeruginosa* functional evidence |
| PA3408 | Q9HYJ7 | HasR | Hemophore-dependent heme receptor and signaling receptor in the Has system; functionally distinct from PhuR. | PMID: 25616666; 31390127 | *P. aeruginosa* functional evidence |
| PA3790 | G3XD89 | OprC | TonB-dependent copper importer. Transports Cu(I)/Cu(II) using conserved metal-binding residues; tied to azurin-dependent copper physiology and virulence. | PMID: 8760927; 31790504; 34762655 | *P. aeruginosa* functional and structural evidence |
| PA4710 | Q9HV88 | PhuR | Heme receptor in the Phu heme-acquisition system; major heme uptake route relative to HasR under several conditions. | PMID: 25616666 | *P. aeruginosa* functional evidence |
| PA3268 | Q9HYX3 | ActA / OptR | Alpha-carboxylate siderophore receptor. Transports rhizoferrin-Fe and staphyloferrin A-Fe. | PMID: 37758521 | *P. aeruginosa* functional evidence |
| PA2289 | Q9I1I6 | OptQ | Predicted iron/siderophore TBDR; no experimentally assigned ligand found. | PMID: 29555629 |  |
| PA1922 | Q9I2I2 | CirA-like | Zinc-homeostasis-associated TBDR; also annotated as CirA-like. No separate receptor-specific substrate paper beyond zinc-homeostasis work was found. | PMID: 26290475 | 35% amino-acid identity to *E. coli* CirA P17315. |
| PA1365 | Q9I3X9 | OptN | Predicted alcaligin/IutA-like receptor. Experimental work in the ChtA context did not support a role in tested citrate-hydroxamate siderophore utilization. | PMID: 16549659; 37758521; 29555629 | Alcaligin/IutA-like annotation |
| PA1271 | Q9I473 | BtuB-like | Predicted cobalamin/vitamin B12 receptor by homology to *E. coli* BtuB. No *P. aeruginosa* receptor-specific experimental paper demonstrating B12 uptake was found. | PMID: 29555629 | *E. coli* BtuB-like homology; 29% amino-acid identity to *E. coli* BtuB P06129. |
| PA0781 | Q9I5F7 | ZnuD-like | Zinc-restriction-inducible TonB-dependent outer-membrane protein; contributes to Zn entry with PA1922, PA2911 and PA4837/CntO. | PMID: 26290475 | *P. aeruginosa* zinc-homeostasis evidence |

PMIDs are provided for the primary or most directly relevant literature supporting each assignment. Rows marked as predicted/unresolved indicate that no receptor-specific substrate paper was identified.

Abbreviations: CSS, cell-surface signaling; TBDR/TBDT, TonB-dependent receptor/transporter.

Supplemental table 2: TBDRs in *P. aeruginosa* with experimental structures available

| Pseudomonas. com accession | PDB accession |
| --- | --- |
| PA2398 | 1XKH |
| PA4221 | 1XKW |
| PA2466 | 8B43 |
| PA4514 | 5FOK |
| PA0931 | 5FP2 |
| PA2688 | 5MZS |
| PA3790 | 6Z8Y |
| PA4710 | 8THE |
| PA1910 | 8S34, 9EX3, 9F2T, 9FVQ |
| PA3063 | 9H80 |
| PA3648 | 9PXG |

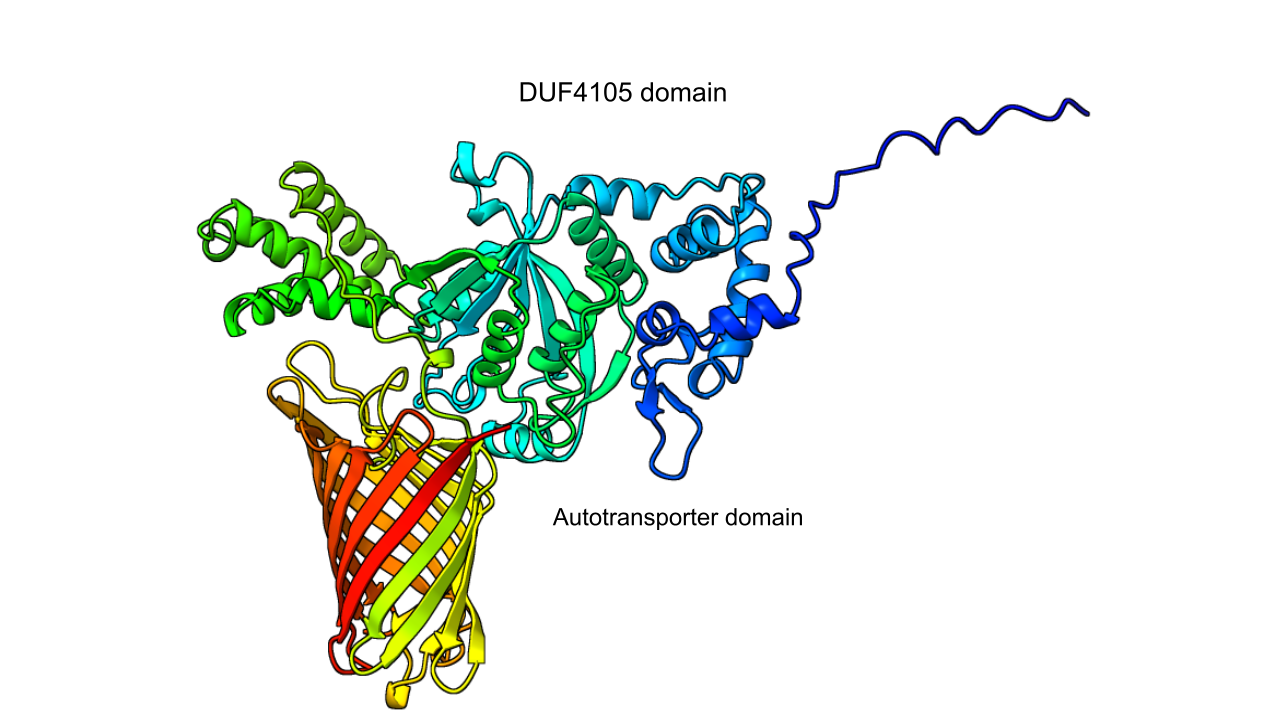

**Supplementary Figure 2. AlphaFold 3 model of PA4579.** Cartoon representation of the predicted structure of PA4579 (UniProt Q9HVK5), showing the N-terminal DUF4105 passenger domain and C-terminal 12-stranded autotransporter β-barrel domain. The model is coloured from the N-terminus (blue) to the C-terminus (red).

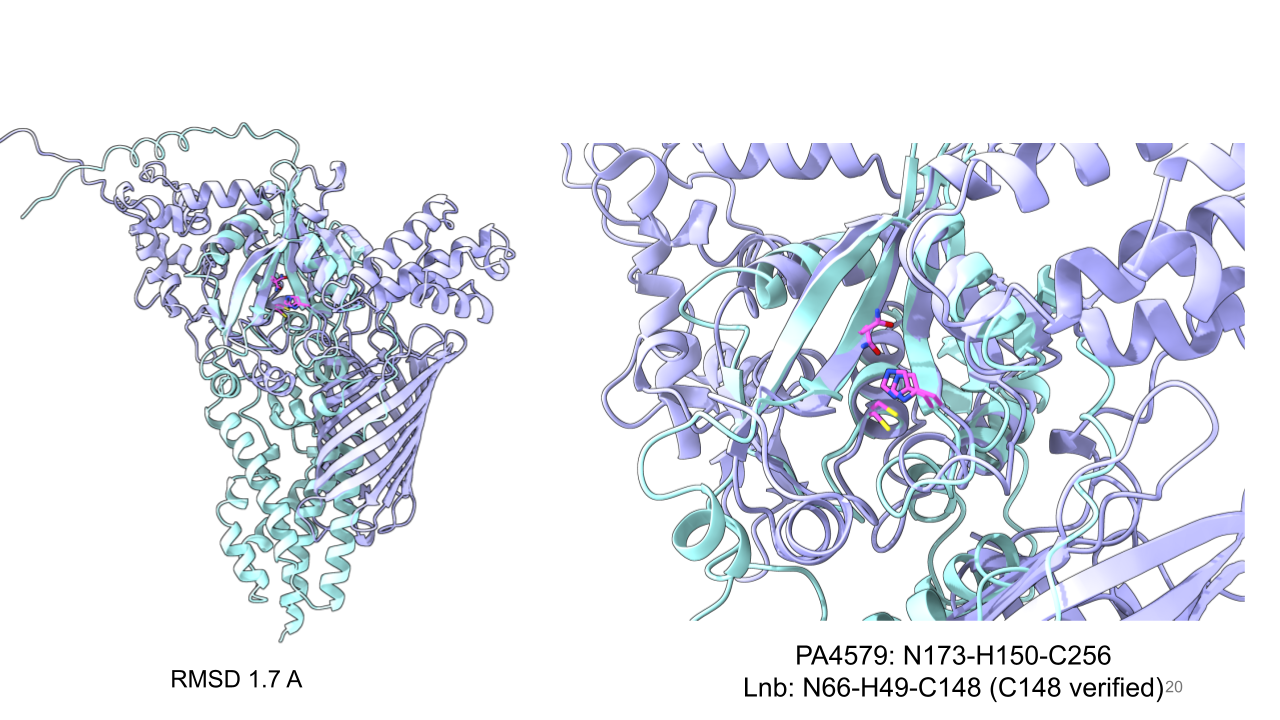

**Supplementary Figure 3. PA4579-1B Structural similarity between PA4579 and the *Bacteroides* lipoprotein *N*-acyltransferase Lnb.** Left, superposition of the AlphaFold 3 models of PA4579 (light purple) and Lnb (cyan). Following distance-based pruning, the conserved DUF4105 structural cores aligned with an RMSD of approximately 1.7 Å across 114 Cα atom pairs. Right, close-up view showing the structural correspondence between the putative catalytic residues Asn173, His150, and Cys256 of PA4579 and Asn66, His49, and Cys148 of Lnb. The functional importance of Lnb Cys148 has been demonstrated experimentally, whereas the roles of Lnb Asn66 and His49 remain structurally inferred.

Supplemental table 3: The list of primers for cloning

| **Name** | **Sequence (5' → 3')** |
| --- | --- |
| PA0151_Fwd | CGAAAACCTGTATTTTCAGGGCGCCGCCGAGGCGGTTCGCTAC |
| PA0165_Fwd | CGAAAACCTGTATTTTCAGGGCGCCGAAGACCATGACATGACGCCG |
| PA0192_Fwd | CGAAAACCTGTATTTTCAGGGCGCCGCCGATGCTCCCGCG |
| PA0234_Fwd | CGAAAACCTGTATTTTCAGGGCGCCCTGGAATGGATGAGCAACAGCGTC |
| PA0328_Fwd | CGAAAACCTGTATTTTCAGGGCGCCTATCAATACGGGGAATACGCC |
| PA0434_Fwd | CGAAAACCTGTATTTTCAGGGCGCCGATGAGGAACCCACCGAACTGCCC |
| PA0470_Fwd | CGAAAACCTGTATTTTCAGGGCGCCGCCCCGGCCGAACAG |
| PA0696_Fwd | CGAAAACCTGTATTTTCAGGGCGCCCAGCCGCCGGCGCCC |
| PA0781_Fwd | CGAAAACCTGTATTTTCAGGGCGCCGCCGACGGCCGCCCC |
| PA0994_Fwd | CGAAAACCTGTATTTTCAGGGCGCCGTGGAATTCAACGCCGAGTTCCTGAACAAC |
| PA1106_Fwd | CGAAAACCTGTATTTTCAGGGCGCCCAGGTGATGGCCCGC |
| PA1271_Fwd | CGAAAACCTGTATTTTCAGGGCGCCGAACCGGTCAGCCTTGCCGAC |
| PA1302_Fwd | CGAAAACCTGTATTTTCAGGGCGCCGCCGAACGACGCTTCGACCTG |
| PA1322_Fwd | CGAAAACCTGTATTTTCAGGGCGCCTGCTCCTCGCCGGCCC |
| PA1343_Fwd | CGAAAACCTGTATTTTCAGGGCGCCGCCGACGACGGCGAC |
| PA1365_Fwd | CGAAAACCTGTATTTTCAGGGCGCCCAGGAAAAGCTCGACGTGGAT |
| PA1613_Fwd | CGAAAACCTGTATTTTCAGGGCGCCGACCTGTTCAACGATCCC |
| PA1763_Fwd | CGAAAACCTGTATTTTCAGGGCGCCGAGGAAGCCTCGGTCGAC |
| PA1910_Fwd | CGAAAACCTGTATTTTCAGGGCGCCGCCGAGGCGCCCGCC |
| PA1922_Fwd | CGAAAACCTGTATTTTCAGGGCGCCGCGCCGGTCGACCTGGAG |
| PA1951_Fwd | CGAAAACCTGTATTTTCAGGGCGCCGCGACGGAATCCGAGGTC |
| PA1974_Fwd | CGAAAACCTGTATTTTCAGGGCGCCGATATCCTGCTGTACGACAAGGAC |
| PA2057_Fwd | CGAAAACCTGTATTTTCAGGGCGCCCAGCAGGAAGGGCCGTCG |
| PA2064_Fwd | CGAAAACCTGTATTTTCAGGGCGCCGCGCAGGACAATGGCATG |
| PA2070_Fwd | CGAAAACCTGTATTTTCAGGGCGCCGCGGAGGAACAGGCGCCT |
| PA2089_Fwd | CGAAAACCTGTATTTTCAGGGCGCCTGCCTTGGCCAGCCCGGC |
| PA2289_Fwd | CGAAAACCTGTATTTTCAGGGCGCCTGCGCCGTCCAGGGC |
| PA2335_Fwd | CGAAAACCTGTATTTTCAGGGCGCCGACGACACCCCCGCGG |
| PA2421_Fwd | CGAAAACCTGTATTTTCAGGGCGCCGACGCCGTCGCGCTG |
| PA2474_Fwd | CGAAAACCTGTATTTTCAGGGCGCCCTGCCCGGCACGCTG |
| PA2590_Fwd | CGAAAACCTGTATTTTCAGGGCGCCTGCGCGCTGCAGGGC |
| PA2911_Fwd | CGAAAACCTGTATTTTCAGGGCGCCGAAACCGAGGAAGCCGCCCTC |
| PA3063_Fwd | CGAAAACCTGTATTTTCAGGGCGCCATGGCCAACTCGTCTGCC |
| PA3268_Fwd | CGAAAACCTGTATTTTCAGGGCGCCGAAGAGGAAGCCAAGGAG |
| PA3408_Fwd | CGAAAACCTGTATTTTCAGGGCGCCCAAGACGGGGCGGACTCG |
| PA3422_Fwd | CGAAAACCTGTATTTTCAGGGCGCCTGCGCCACCCAGGCC |
| PA3497_Fwd | CGAAAACCTGTATTTTCAGGGCGCCGCTTCGCCGCCGTCC |
| PA3535_Fwd | CGAAAACCTGTATTTTCAGGGCGCCGCCTATGTCGAAGCCGGT |
| PA3772_Fwd | CGAAAACCTGTATTTTCAGGGCGCCGCCGACCTCAACGCC |
| PA3901_Fwd | CGAAAACCTGTATTTTCAGGGCGCCGCCGAGATCCGCACCGAT |
| PA3923_Fwd | CGAAAACCTGTATTTTCAGGGCGCCGTGACCTTCAACATCGGGGAA |
| PA4084_Fwd | CGAAAACCTGTATTTTCAGGGCGCCGTCATCGAGCCAGGCCGCTAT |
| PA4156_Fwd | CGAAAACCTGTATTTTCAGGGCGCCCGGGAAAACCCGCCCG |
| PA4168_Fwd | CGAAAACCTGTATTTTCAGGGCGCCGCTCCCATCGCCCTTGAC |
| PA4390_Fwd | CGAAAACCTGTATTTTCAGGGCGCCGATACGGTCTGGATGAAGAACG |
| PA4391_Fwd | CGAAAACCTGTATTTTCAGGGCGCCGACACCGTATGGCTGGAC |
| PA4579_Fwd | CGAAAACCTGTATTTTCAGGGCGCCGCGCCCACGGTTCCC |
| PA4652_Fwd | CGAAAACCTGTATTTTCAGGGCGCCGCGGACTACCTGGCC |
| PA4675_Fwd | CGAAAACCTGTATTTTCAGGGCGCCGCCGATCCGGTGGAGCAG |
| PA4710_Fwd | CGAAAACCTGTATTTTCAGGGCGCCGGGAACGCCGTCCCG |
| PA4824_Fwd | CGAAAACCTGTATTTTCAGGGCGCCATGACCCTGCACGACCGG |
| PA4837_Fwd | CGAAAACCTGTATTTTCAGGGCGCCTGCAGCAGTCCGGCGCTCTGG |
| PA4897_Fwd | CGAAAACCTGTATTTTCAGGGCGCCAGCGTCGGCCAGCACAG |
| PA5112_Fwd | CGAAAACCTGTATTTTCAGGGCGCCGCTCCTTCGCCCTATTCG |
| PA5325_Fwd | CGAAAACCTGTATTTTCAGGGCGCCTGCACCCACGCCAGC |
| PA0151_Rev | AGCTTGTCGACGGAGCTCGAATTCGTCAGAACTCCACCGTCGTCGACA |
| PA0165_Rev | AGCTTGTCGACGGAGCTCGAATTCGTCAGAAGTGGTACTTGACCAG |
| PA0192_Rev | AGCTTGTCGACGGAGCTCGAATTCGTCAGAAGTCGTAGCGCAGCGATAC |
| PA0234_Rev | AGCTTGTCGACGGAGCTCGAATTCGTCAGAAGGTGAATTCCATATTCAACGT |
| PA0328_Rev | AGCTTGTCGACGGAGCTCGAATTCGTCAGAACTGCCAGTTCAC |
| PA0434_Rev | AGCTTGTCGACGGAGCTCGAATTCGTCAGAAGCGCGTCTGCACGCTCAG |
| PA0470_Rev | AGCTTGTCGACGGAGCTCGAATTCGTCACCACTTGTAGCTGACGCT |
| PA0696_Rev | AGCTTGTCGACGGAGCTCGAATTCGTCAGAAGCGCGTGTTGACCTCGAG |
| PA0781_Rev | AGCTTGTCGACGGAGCTCGAATTCGTCAGAACGCCACCTTCACCCCCGC |
| PA0994_Rev | AGCTTGTCGACGGAGCTCGAATTCGTCAGATCGAGGACCGACGGCAACTGAG |
| PA1271_Rev | AGCTTGTCGACGGAGCTCGAATTCGTCACAGAGCCTGGTAGCGCACGCT |
| PA1302_Rev | AGCTTGTCGACGGAGCTCGAATTCGTCAGAAGAACTGGGTGACGCTGATCTTGGC |
| PA1322_Rev | AGCTTGTCGACGGAGCTCGAATTCGTCAGAACGCGTAGATCGCCGAAAGAAT |
| PA1343_Rev | AGCTTGTCGACGGAGCTCGAATTCGTCAGAGACGCAGGCCGAG |
| PA1365_Rev | AGCTTGTCGACGGAGCTCGAATTCGTCACCAGTTGTGCGTATAGCT |
| PA1613_Rev | AGCTTGTCGACGGAGCTCGAATTCGCTAGAACGTCACCTCGGC |
| PA1763_Rev | AGCTTGTCGACGGAGCTCGAATTCGTTACTGGGCCTTCAGCCCTTT |
| PA1910_Rev | AGCTTGTCGACGGAGCTCGAATTCGTCAGAAGTCCGCCGTGAGGGAAAGGGA |
| PA1922_Rev | AGCTTGTCGACGGAGCTCGAATTCGTCAGAAGCTCGCCACCAGGCCG |
| PA1951_Rev | AGCTTGTCGACGGAGCTCGAATTCGTCAGAAGTAGTAGGGGAATTTCACGCC |
| PA1974_Rev | AGCTTGTCGACGGAGCTCGAATTCGTCACCAGTTCAGTACGGC |
| PA2057_Rev | AGCTTGTCGACGGAGCTCGAATTCGTCAACCACCGAAGTGATAGCTGAAGTC |
| PA2064_Rev | AGCTTGTCGACGGAGCTCGAATTCGTCAGAACCACATGCGGATACC |
| PA2070_Rev | AGCTTGTCGACGGAGCTCGAATTCGTCAGAAGCTGTAGCTGAGGCGGC |
| PA2089_Rev | AGCTTGTCGACGGAGCTCGAATTCGTTAGAAGGTGTAGGCCGCGCCGAG |
| PA2289_Rev | AGCTTGTCGACGGAGCTCGAATTCGTCAGAACTGGTATTCGACCCCG |
| PA2335_Rev | AGCTTGTCGACGGAGCTCGAATTCGTCAGAAGTCGTAGCGGGCGGT |
| PA2421_Rev | AGCTTGTCGACGGAGCTCGAATTCGTCAGAACGCCAGCAGGTA |
| PA2474_Rev | AGCTTGTCGACGGAGCTCGAATTCGTCAGAAGACCTGCGAGTAGCG |
| PA2590_Rev | AGCTTGTCGACGGAGCTCGAATTCGCTAGAAGCGGTATGCCGCGCG |
| PA2911_Rev | AGCTTGTCGACGGAGCTCGAATTCGTCAGAAACTGAAGTCGGCGGTGACGTT |
| PA3063_Rev | AGCTTGTCGACGGAGCTCGAATTCGTCAGCGGCCGAAGCG |
| PA3268_Rev | AGCTTGTCGACGGAGCTCGAATTCGTCAGAAACCGACGCT |
| PA3408_Rev | AGCTTGTCGACGGAGCTCGAATTCGTCAGAACTGGTATTCGAGGGTGCCTTG |
| PA3422_Rev | AGCTTGTCGACGGAGCTCGAATTCGTCAGAAGTTCACCCCGAAGCT |
| PA3497_Rev | AGCTTGTCGACGGAGCTCGAATTCGTCAGAAGTAGTAGCCGATGTTCAGGTT |
| PA3535_Rev | AGCTTGTCGACGGAGCTCGAATTCGTCAGAAACGCCAGTCGACGGC |
| PA3772_Rev | AGCTTGTCGACGGAGCTCGAATTCGTCAGAACACATGGACCAGGCG |
| PA3901_Rev | AGCTTGTCGACGGAGCTCGAATTCGTCAGAAACCGACCGAGGCCTGTAC |
| PA3923_Rev | AGCTTGTCGACGGAGCTCGAATTCGTCAGAAGTTCATGCCGAAGCT |
| PA4084_Rev | AGCTTGTCGACGGAGCTCGAATTCGTCACAGCGTGCGCTCCCGGTC |
| PA4156_Rev | AGCTTGTCGACGGAGCTCGAATTCGCTAGAAGGTGCCCTTGATGCC |
| PA4168_Rev | AGCTTGTCGACGGAGCTCGAATTCGTCAGAGCGAGTACTTCACCGTGAACAT |
| PA4390_Rev | AGCTTGTCGACGGAGCTCGAATTCGTTACCAGCCGACCCCGAG |
| PA4391_Rev | AGCTTGTCGACGGAGCTCGAATTCGTCACCAGCCGACGCC |
| PA4579_Rev | AGCTTGTCGACGGAGCTCGAATTCGTCAGTAGTAGTACCAGCGCAG |
| PA4652_Rev | AGCTTGTCGACGGAGCTCGAATTCGTCATGGCGTATCTCCGAT |
| PA4675_Rev | AGCTTGTCGACGGAGCTCGAATTCGTCAGAACTCCACGCTATACGTCAG |
| PA4710_Rev | AGCTTGTCGACGGAGCTCGAATTCGTCAGATGTCCCAGACCAGGTTG |
| PA4824_Rev | AGCTTGTCGACGGAGCTCGAATTCGTCAGAAACGCAAGGTCGCTCC |
| PA4837_Rev | AGCTTGTCGACGGAGCTCGAATTCGTCAGTAGTTCAGGGTCAGGCTCATGGTCA |
| PA4897_Rev | AGCTTGTCGACGGAGCTCGAATTCGTCAGAACTTCGCCTGCATCCC |
| PA5112_Rev | AGCTTGTCGACGGAGCTCGAATTCGTCAGAAGTCCAGGCTCAG |
| PA5325_Rev | AGCTTGTCGACGGAGCTCGAATTCGTCAGAGTTCGAACGGAATGGT |
| PA1106_Rev | AGCTTGTCGACGGAGCTCGAATTCGTCAGAAGAACGGACGCGA |

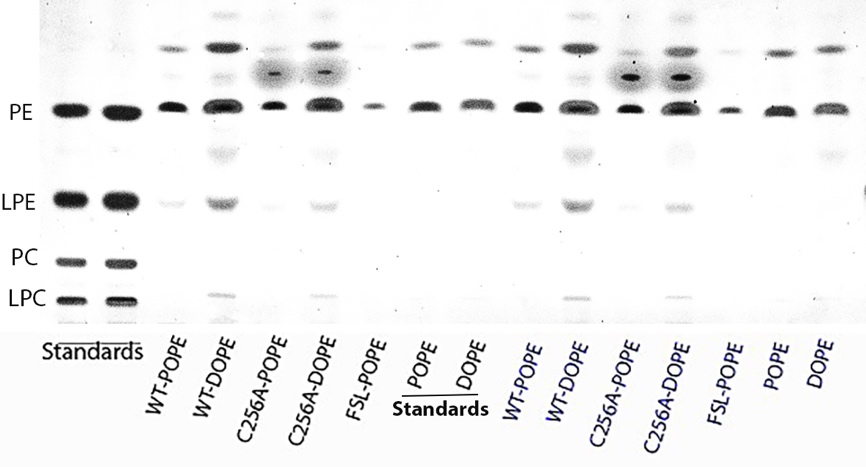

**Supplementary Figure 4. Uncropped plate for main Figure 11.**

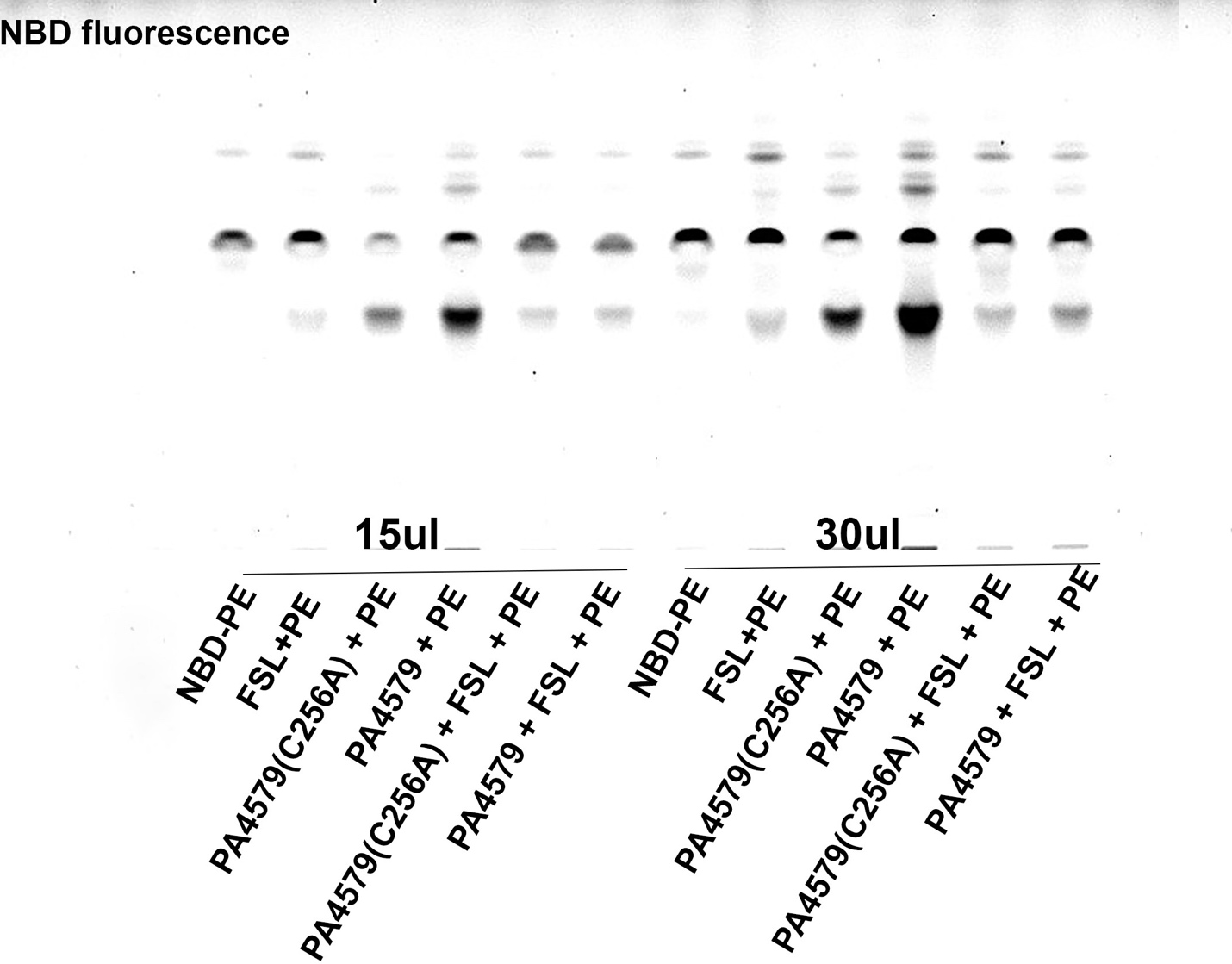

**Supplementary Figure 5. Uncropped plate for main Figure 12-A.**

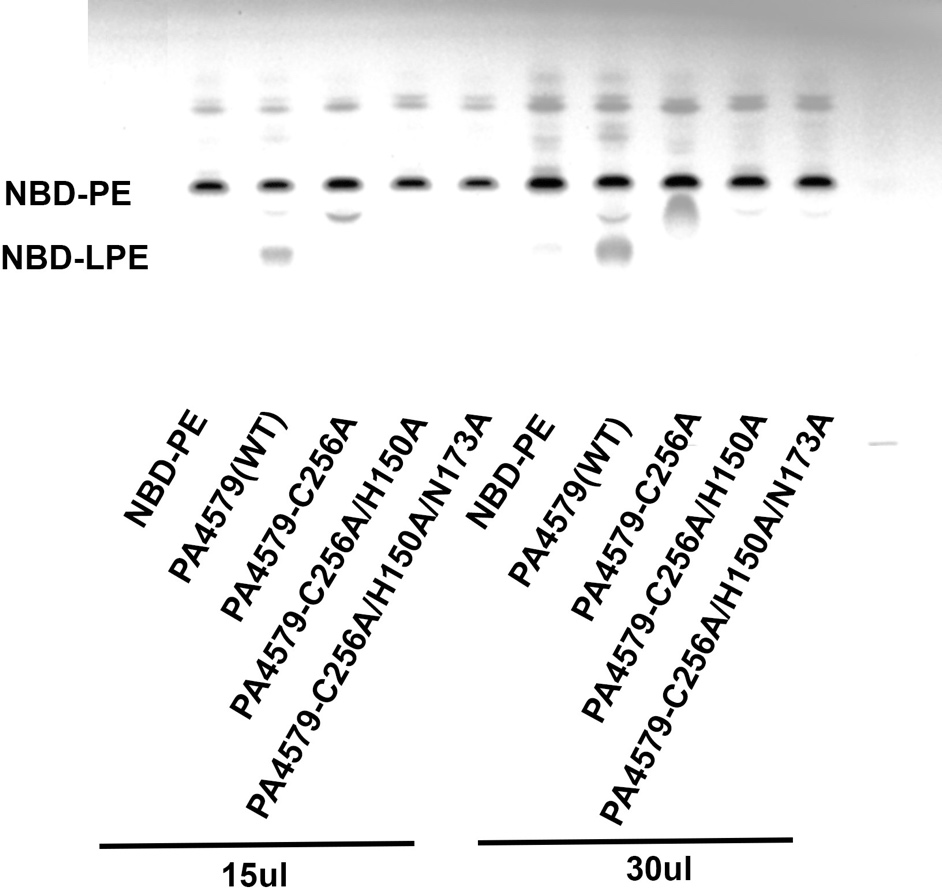

**Supplementary Figure 6. Uncropped plate for main Figure 12-B.**
